# Pain Matters: Nociceptive signaling drives neural encoding of inflammatory responses

**DOI:** 10.64898/2026.11.02.763662

**Authors:** Megan Sammons, Rani Assa Polansky, Itay Zayalat, Samantha Breslauer Morris, Liwen Deng, Hilla Azulay-Debby, Neomi Singer, Isaac Chiu, Asya Rolls

## Abstract

Pain is a cardinal sign of inflammation, constructed by the brain from peripheral nociceptive input. Yet, its role in the brain’s response to inflammation remains unknown. Here, we demonstrate that nociceptive signaling is essential for the brain to represent and replay inflammatory states. Analysis of human fMRI and mouse activity mapping revealed that peripheral inflammation generates a distinct brain state. In mice, ablation of TRPV1⁺ sensory neurons or pharmacological analgesia disrupted this organization and prevented the insular cortex from forming retrievable immune representations. Conversely, augmenting nociceptive signaling during inflammation strengthened subsequent immune recall. These findings identify nociceptive input as a critical signal in brain–immune dialogue, enabling the encoding and retrieval of inflammatory events.

## Main Text

Pain is an unpleasant sensation that alerts the organism to potential injury and drives protective behavioral reflexes^1^. Nociceptive pain is important for rapid detection of external threats, eliciting withdrawal, avoidance and other behaviors that limit damage. Pain also accompanies many infectious and inflammatory conditions^2^. In these contexts, immediate behavioral avoidance may be less relevant and other sickness-related behaviors such as fatigue, reduced feeding, and social withdrawal, may be more adaptive. This raises an important question: what advantage does pain provide during inflammation?

Indeed, recent studies have shown that nociceptive neurons actively regulate local immune responses by releasing neuropeptides and other mediators from peripheral nerve terminals^3,4^. However, ultimately, nociceptive signals are relayed to the central nervous system, where they are integrated into distributed brain activity^5,6^ and reorganize activity across brain networks^7–9^. Therefore, we reasoned that nociceptive signaling participates in how the brain processes information and represents the inflammatory state.

Nociceptive sensory neurons provide temporally precise and anatomically localized information about tissue damage^10^, which may enable the brain to encode the location and nature of inflammation. We know that brain activity changes during inflammation, and we and others have shown that the brain forms representations of inflammatory states, “immune engrams”, particularly in the insular cortex^11,12^. Reactivation of these engrams can replay the inflammatory state, much like other forms of memory^13^. Since not all experiences are encoded into lasting memories, salience is required to mark significant events. We hypothesized that nociceptive signals during inflammation serve this role, acting as a salience signal that supports the organization of neural activity and the formation of a retrievable representation of immune states.

Using reanalysis of human fMRI data and activity mapping in mice, we demonstrate that peripheral inflammation generates a distinct brain-wide network state and that nociceptive signaling is required for both this state and immune engram formation. Disrupting nociceptive input, through TRPV1⁺ neuron ablation or pharmacological analgesia, abolishes retrievable immune encoding, while augmenting nociceptive signaling strengthens immune recall. Together, these findings recast pain not merely as a symptom of inflammation, but as a critical instructive signal that enables the brain to remember it.

### Inflammation induces an “inflammatory network state” in humans and mice

Prior neuroimaging work has shown that inflammation alters functional connectivity in specific brain regions and networks, including the brainstem, insula, thalamus, default mode, and salience networks^14,15^. Whether these are isolated effects or part of a broader whole-brain state that organizes a coherent systemic response has not been tested directly. To address this question, we used an existing human dataset with functional MRIs (fMRIs) acquired before (Baseline) and throughout (Onset, Acute, and Peak) experimental endotoxin administration, a model of LPS-induced systemic inflammation^16^. Although the dataset is small (n = 7), its longitudinal design enabled us to track within-subject connectivity changes as inflammation progressed (Fig. 1A). Brain-wide activity was quantified using a 116-region parcellation spanning seven cortical networks and 16 subcortical structures, including the amygdala, thalamus, and striatum. These regions are both immunologically relevant and typically missing from cortex-only parcellations (Fig. S1A)^17–19^.

**Fig. 1.**
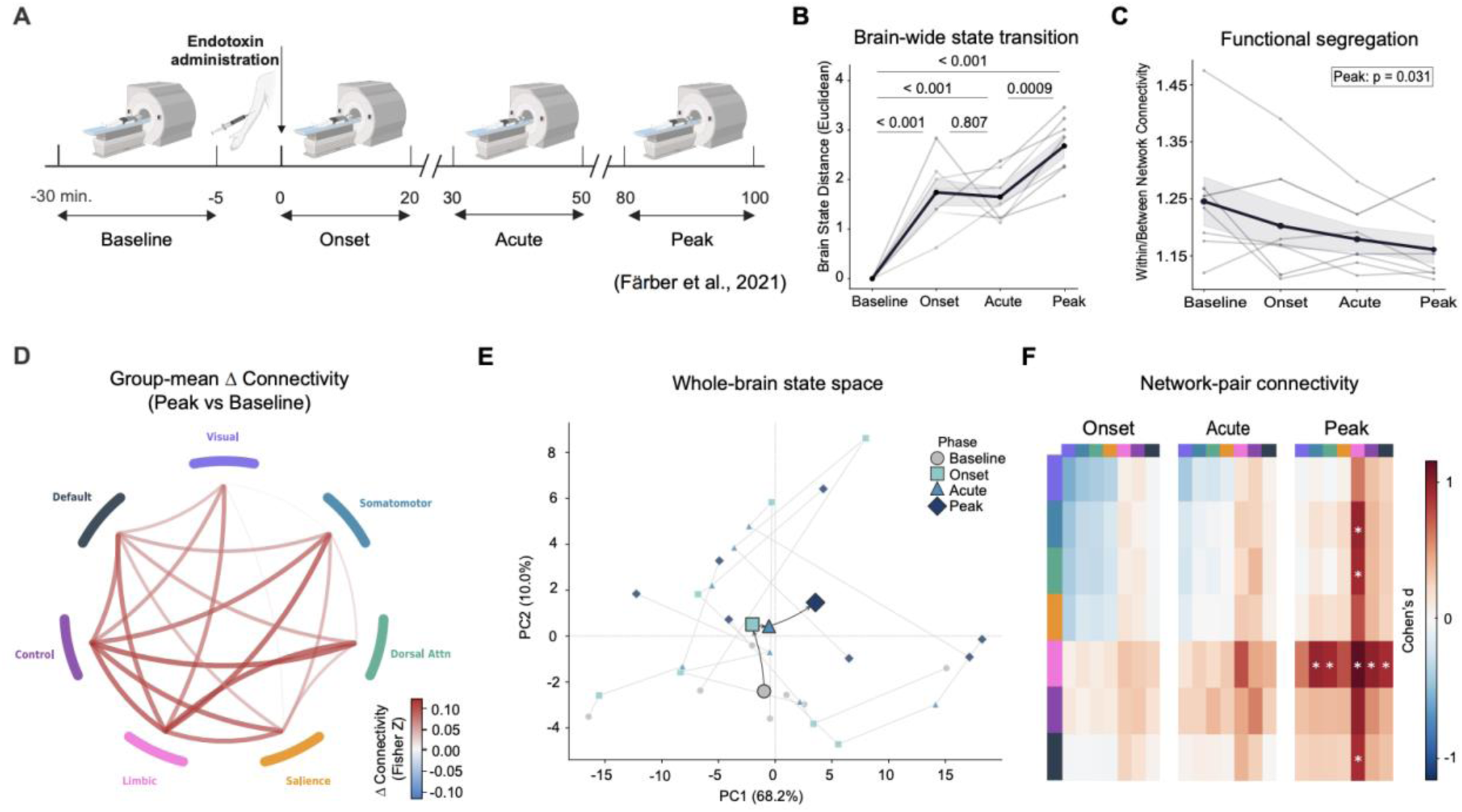
Acute endotoxemia in humans induces a coordinated and structured brain state. Systemic inflammation triggers a rapid, progressive reorganization of the human functional connectome, characterized by increasing global state divergence and a transition toward network desegregation. **(A)** Experimental timeline. Healthy subjects (n=7) underwent repeated fMRI scans around a single intravenous endotoxin challenge (t=0), covering four phases: Baseline (−30 to −5 min), Onset (0–20 min), Acute (30–50 min), and Peak (80–100 min). **(B)** Brain-wide state transition. Euclidean distance of each subject’s whole-brain connectivity vector from their Baseline, computed across 116 parcels. Distance increased significantly at every post-injection phase (one-sample t-test vs. zero: Onset t(6) = 6.54, p = 0.0006, d = 2.47; Acute t(6) = 8.52, p = 0.0001, d = 3.22; Peak t(6) = 11.17, p < 0.0001, d = 4.22). The shift plateaued between Onset and Acute (paired t-test: p = 0.81) before escalating further at Peak (Acute vs. Peak: p = 0.0009). Black line, group mean ± 95% CI; gray lines, individual subjects. **(C)** Functional segregation. Within-to between-network connectivity ratio across phases. Segregation was lower at Peak than Baseline (one-sided paired t-test: t(6) = 2.28, p = 0.031, d = −0.86). **(D)** Chord diagram of mean Δ functional connectivity (Fisher Z) networks (Limbic: paired t-test: t(6) = 3.46, p = 0.014, d = 1.31, no network survived FDR correction). Edge thickness reflects effect magnitude; red edges denote increased coupling. **(E)** Whole-brain connectivity state space. PCA of all 116 × 116 connectivity matrices (PC1 = 68.2%, PC2 = 10.0%). Each scan is a point, colored and shaped by phase. Subjects trace a directed trajectory away from Baseline (gray circles) toward Peak (dark blue diamonds) across phases. Network-level connectivity change (Peak − Baseline). **(F)** Pairwise network connectivity. Cohen’s d heatmaps for each network pair relative to Baseline at Onset, Acute, and Peak. No network pairs survived FDR correction: asterisks mark uncorrected p-values. *All plots: n=7*.

For each subject, we first quantified how much their connectivity changed from baseline (p < 0.001 at Onset, Acute, and Peak) (Fig. 1B). Distance increased at every timepoint relative to Baseline and scaled with inflammatory progression (Peak-Acute: p = 0.0009). Leave-one-out sensitivity analyses confirmed that these effects did not depend on any single subject (p < 0.001) (Fig. S1). Thus, endotoxin drives a reliable, directed shift in whole-brain functional architecture that accelerates as inflammation progresses.

To characterize the direction of this network shift, we quantified functional segregation, the relative strength of within-network versus between-network connectivity (Fig. 1C). At Baseline, within-network connections were stronger, consistent with canonical resting-state modularity^20^. This ratio progressively declined, reflecting a reduction in network segregation as the brain converged toward a less modular brain state at Peak (p = 0.031). This means that inflammation progressively erodes the brain’s default compartmentalized organization, shifting it toward a more globally integrated state. This trajectory was consistent across subjects. Principal component analysis (PCA) of all connectivity matrices showed individuals moving along a shared path from Baseline toward Peak in brain state space (Fig. 1D). Connectivity changes were not uniform across networks. Peak-to-Baseline differences were driven specifically by the Limbic network (p = 0.014), while no other network showed significant changes (all p > 0.15) (Fig. 1E-F). This aligns with sickness behaviors that accompany systemic inflammation, such as fatigue, reduced feeding, and social withdrawal, coordinated by limbic circuitry^21^. Importantly, these changes extend beyond canonical pain-processing regions. We compared the spatial distribution of fMRI activation across all time points against the Neurological Pain Signature (NPS) across brain parcels and found no significant overlap (p = 0.83) (Fig. S1I) ^9^. This indicates that this reorganization reflects a broader brain-wide response rather than selective engagement of pain circuits.

To ask whether nociceptive input drives this reorganization, we turned to mice, where nociceptive neurons can be pharmacologically targeted. We first determined whether an inflammatory network state is conserved across species. We used Zymosan-induced peritonitis (ZIP), a model of sterile peritoneal inflammation, and analyzed changes in brain activity using TRAP2-Ai14D mice. This transgenic line expresses a tamoxifen-inducible Cre under the activity-dependent *Fos* promoter in combination with the Ai14D-tdTomato reporter, allowing long-term labeling of activated neurons at a specific time point. We utilized this system to label active neurons (TRAP) during the inflammatory window. Whereas the human analysis measured functional connectivity across 116 parcellated regions spanning functional networks, the mouse whole-brain data were organized across anatomically defined regions using hierarchical co-activation clustering. In agreement with the human data, the inflamed mouse brain showed a similar pattern of increased coupling across regions and a shift toward a more integrated network state (p < 0.0001) (Fig. S2), driven by sensory relay regions.

Together, these results demonstrate that systemic inflammation induces a coordinated shift in brain network architecture, characterized by reduced segregation and increased cross-network integration. Rather than selectively activating pain-related circuits, inflammation drives a global reorganization of brain function. Having established conservation, we could now exploit mouse genetic and pharmacologic tools to mechanistically test the role of nociceptive input to organize this brain-wide state.

### TRPV1⁺ sensory neurons innervate the peritoneal cavity

The parietal peritoneum, the site of the Zymosan-induced peritonitis, receives significant sensory innervation by thoracic DRG neurons from T6–T12^22^. Nociceptors are a major class of sensory neurons that receive input from the abdominal viscera^23^, though their precise anatomical distribution remains unresolved. We mapped this innervation using viral tracing from the peritoneal wall and dissecting T6–T12 DRGs (Fig. 2A). This revealed dense labeling in thoracic DRGs and co-expression of the nociceptive ion channel TRPV1^+^ in a subset of labeled neurons (Fig. 2B). thus, TRPV1^+^ neurons are a source of sensory input from the peritoneum to the spinal cord.

**Fig. 2.**
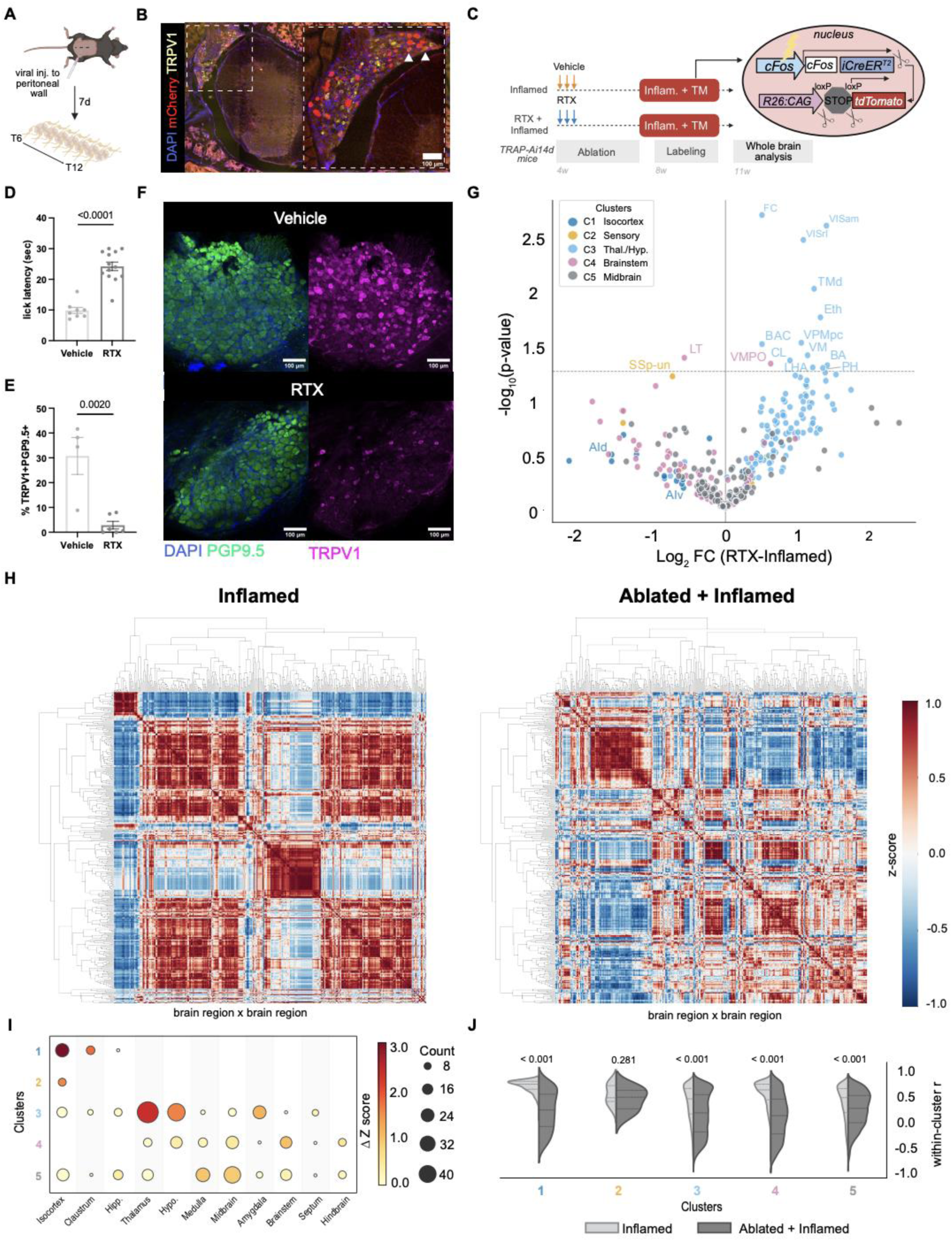
Peripheral TRPV1+ sensory neurons organize the brain-wide inflammatory state. TRPV1+ sensory neurons innervate the peritoneal wall, and their ablation disrupts the systemic integration of inflammatory signals, leading to a collapse of coordinated neural network activity across anatomical clusters. **(A)** Experimental scheme for viral tracing from the peritoneum. Mice were injected with a viral tracer into the peritoneal wall and sections of the spine from T6-T12 were collected 7 days later **(B)** Representative immunofluorescent images taken from cross section of the spinal cord and Dorsal Root Ganglia (DRG) labeled with DAPI (blue), virus (mCherry), and TRPV1 (yellow). **(C)** Experimental workflow for whole-brain activity mapping. TRAP2-Ai14D mice received 3 days of consecutive, increasing resiniferatoxin (RTX) to ablate TRPV1+ neurons or vehicle injection. Following a 4-week recovery, mice were administered TM during ZIP to label active neurons for whole-brain quantification and network analysis. **(D)** Behavioral validation of nociceptor ablation by hot plate assay. Quantification of response latency after being placed on hot plate at 57C for up to 30 seconds (p < 0.0001, unpaired t-test, vehicle: n = 7; RTX: n = 13). **(E)** Quantification of the percent of TRPV1+ PGP9.5+ neurons in the DRG (p = 0.0020, unpaired t-test, vehicle: n = 4; RTX: n = 6). **(F)** Representative immunofluorescent images of taken from T6-T12 DRGs stained for DAPI (blue), PGP9.5 (green), and TRPV1 (magenta). **(G)** Volcano plot showing the relationship between magnitude of change (log2 fold change) and statistical significance (-log10 p-value) for 361 Allen CCF brain regions between Inflamed and Ablated + Inflamed (RTX). Each dot represents a specific brain region, color coded by Cluster identify from hierarchical clustering. SSp-un: Primary somatosensory, undefined; LT: Lamina Terminalis; VMPO: Ventromedial Preoptic Nucleus; LHA: Lateral Hypothalamic Area; PH: Posterior Hypothalamic Nucleus; BA: Bed Nucleus of the Accessory Olfactory Tract; VM: Ventromedial Nucleus of the Thalamus; VPMpc: Ventral Posteromedial Nucleus of the Thalamus, parvicellular part; CL: Central Lateral Nucleus of the Thalamus; BAC: Bed Nucleus of the Anterior Commissure; Eth: Ethmoid Nucleus of the Thalamus; TMd: Tuberomammillary Nucleus, dorsal part; VISrl: Rostrolateral Visual Area; VISam: Anteromedial Visual Area; FC: Fasciola Cinerea. **(H)** Hierarchical clustering of pairwise regional activity correlations, z-score of Pearson’s r, average linkage used for clustering, permutations = 5000, Inflamed: n=8, Ablated + Inflamed: n=9. **(I)** Bubble plot showing the enrichment of specific anatomical structures across the five functional clusters. Circle size represents region count, and color intensity represents the connectivity loss measured by z-score. **(J)** Split violin plot shows the distribution of within-cluster correlation coefficients for Inflamed vs. Ablated + Inflamed, Cluster 1: p < 0.0001; Cluster 2: p = 0.281; Cluster 3: p < 0.0001; Cluster 4: p < 0.0001; Cluster 5: p < 0.0001, Mann-Whitney U test. *Data are presented as mean ± SEM*.

### Peripheral TRPV1+ sensory neurons are required to organize the inflammatory brain state

TRPV1^+^ is a nociceptive ion channel expressed by peptidergic C-fiber neurons that detect noxious heat and capsaicin and can be sensitized by immune mediators to drive inflammatory pain^24,25,26^. TRPV1-expressing neurons are known to relay peripheral inflammatory signals to the central nervous system^24,27,28^, placing these neurons at the critical boundary between peripheral immune sensing and central processing. Having established TRPV1^+^ neurons as an afferent input from the peritoneum, we next tested whether they are required for the brain-wide activity changes we observed during inflammation.

We tested this by selectively ablating TRPV1⁺ sensory neurons in TRAP2-Ai14D mice (Fig. 2C)^29^. To eliminate TRPV1⁺ neurons, we used Resiniferatoxin (RTX), a high affinity TRPV1⁺ ligand that induces chemical ablation of nociceptors^30^. Two groups of FosTRAP2 mice received either RTX (“Ablated”) or vehicle (“Control”) at 4 weeks old and were allowed to recover for 4 weeks. RTX administration effectively ablated TRPV1⁺ neurons, as confirmed by behavioral assay and histological analysis of DRG neurons (Fig. 2D–E; Fig. S3). Thus, we could ask whether removing nociceptive input changed which brain regions were recruited during inflammation.

Nociceptor ablation shifted the regional pattern of TRAPed neuron density. Thalamic and hypothalamic regions showed increased activation relative to inflamed controls (Fig. 2G), while total brain-wide TRAPed neurons did not differ between groups (p = 0.6825) (Fig. S4A). This suggested that neural activity was instead redistributed across regions.

To determine whether this redistribution reflects how nociceptive signals organize the inflammatory brain state, we performed hierarchical clustering of regional co-activation. Five functional clusters emerged, and their overall architecture differed substantially between TRPV1^+^ ablated and vehicle-treated controls (Fig. 2H-J, S4B). Cluster 1, enriched in isocortical regions including posterior insular, primary somatosensory, motor, and orbitofrontal cortex, lost internal synchrony and collapsed as a network hub in TRPV1-ablated animals (p < 0.001). Cluster 2, enriched in visceral and secondary somatosensory cortex, was the only cluster unaffected by ablation (p = 0.281). Cluster 3, dominated by thalamic and hypothalamic regions, became the most active cluster in ablated animals yet was no longer functionally coupled to the isocortical regions it normally coordinates with (p < 0.001). Clusters 4 and 5, enriched in brainstem and midbrain respectively, also lost synchronization (p < 0.001). Thus, nociceptor ablation systematically altered brain-wide synchronization.

Between-cluster covariance revealed the consequence of these effects. During inflammation, Cluster 1, enriched in isocortical regions, becomes anti-correlated with Cluster 5, represented mainly in midbrain regions (r = -0.83, p = 0.011), an effect not present in TRPV1^+^ ablated animals (r = -0.45, p = 0.263) (Fig. S4C, D). Indeed, the cortex is known to exert top-down inhibitory control over subcortical circuits during pain^31–33^. Our data suggest that this control requires intact peripheral nociceptive input. Subcortically, Clusters 3 and 5 were positively coupled during inflammation (r = +0.73, p = 0.04) but anti-correlated after ablation (r = -0.82, p = 0.008). Importantly, this data suggests that subcortical structures receive inflammatory input regardless of nociceptive drive. What nociception therefore adds is cortical engagement.

Taken together, these findings reveal a dissociation between detection and organization of inflammation by the brain. TRPV1⁺ sensory neurons are essential for organizing signals into a coordinated, brain-wide network architecture. In their absence, cortical interoceptive regions disengage while subcortical processing nodes become hyperactive. Because coordinated network activity is required for the formation of stable neural representations^34^, these findings raise the possibility that nociceptive input may be required not only for organizing the inflammatory network state, but also for encoding a retrievable neural representation of inflammation.

### TRPV1⁺ neurons are required for insular encoding and retrieval of inflammatory responses

Network enrichment analysis revealed that inflammation engages distributed brain circuits, with the isocortex emerging as significantly enriched in Cluster 1 and disrupted following TRPV1⁺ neuron ablation (p < 0.001). Within this network, the posterior insular cortex (pIC), a key region for representing internal physiological states, has been implicated in the storage and retrieval of immune responses^11^. Furthermore, the insular cortex is also known to be involved in pain processing and perception^35,36^. We therefore asked whether TRPV1⁺ sensory neurons are required for the encoding of inflammation within the pIC. Namely, we wanted to compare the representation formed in the presence and absence of TRPV1-nociceptive signaling.

However, it has been shown that in barrier tissues, nociceptive neurons can also regulate the immune responses^3,4^. Thus, we first determined whether TRPV1⁺ neuron ablation altered the initial inflammatory response to ZIP (Fig. S3A). Nociceptor ablation did not significantly alter the cellular composition in the peritoneal lavage fluid (PLF) 24 hours after ZIP (Fig. S3). Since the key immune parameters were comparable in the control and ablated groups, we next tested whether and how the nociceptive input could affect the formation of a retrievable neural representation. For this purpose, we combined activity-dependent neuronal labeling with chemogenetic reactivation of inflammation-associated neuronal ensembles in the pIC of mice treated with or without RTX (Fig. 3A). As before, two groups of FosTRAP2 mice, which received RTX (“Ablated”) or vehicle (“Control”), were injected with AAV-hSyn-DIO-hM3Dq-mCherry into the pIC. This allowed us to TRAP neurons during inflammation and then control their activity using the activating form of DREADDs (Gq). During ZIP, we administered tamoxifen (TM) to label inflammation-associated ensembles in the ablated and control mice. Then, after recovery from the original inflammation, we administered the DREADD agonist JHU37160 (JHU) and assessed the immune response in the PLF 24 hours later by flow cytometry.

**Fig. 3.**
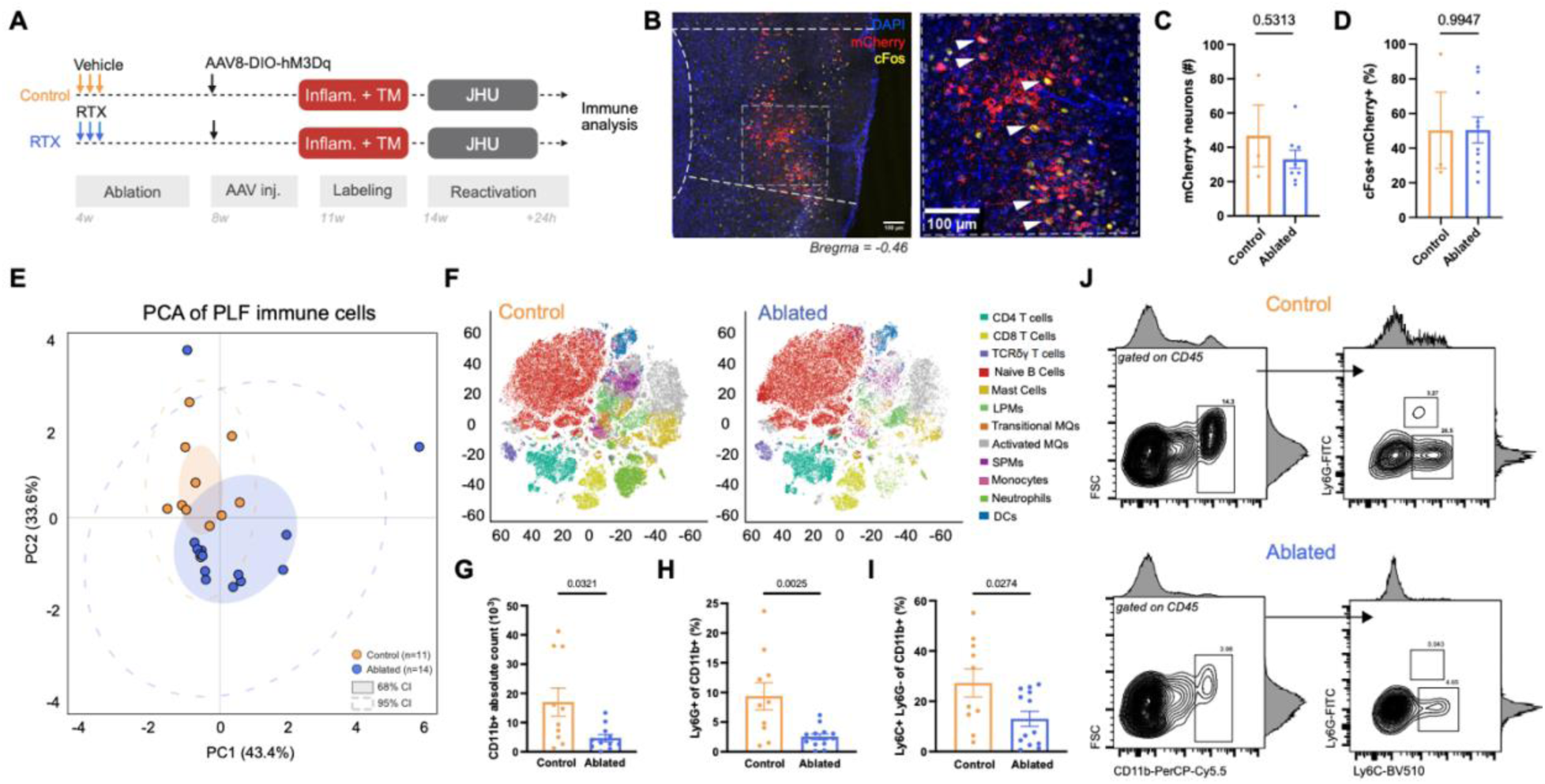
TRPV1+ sensory neurons are required for the insular cortex to drive peripheral immune recruitment. Chemogenetic reactivation of pIC ensembles labeled during inflammation fails to recapitulate peripheral immune response in the absence of peripheral TRPV1+ sensory neurons. **(A)** Experimental timeline for labeling and chemogenetic reactivation of pIC neurons. 4-week-old TRAP2Fos mice received RTX or vehicle for 3 consecutive days. After recovery, the excitatory DREADD, AAV-hSyn-DIO-hM3Dq-mCherry, was injected into the pIC. Then, Tamoxifen (TM) was administered to label active neurons during inflammation. These ensembles were reactivated 3 weeks later using JHU, followed by immune analysis. **(B)** Representative immunofluorescent image of cFos+mCherry+ neurons in the pIC. DAPI (blue), AAV (mCherry), cFos (yellow), scale bar 100um **(C)** Quantification of virally expressed mCherry+ neurons (p = 0.513). Control n = 3, Ablated n = 8. **(D)** Quantification of cFos+mCherry+ neurons in the pIC (p = 0.9947). Control n = 3, Ablated n = 8. **(E)** PCA analysis of myeloid and B cells from the PLF (PERMANOVA p = 0.0012, F = 8.622; PERMDISP p = 0.0615, F = 3.670, 9,999 permutations) **(F)** tSNE embedding of total CD45+ cells from the PLF following reactivation in Control vs Ablated mice. **(G-I)** Bar plots of the total count of PLF CD11b+ cells following reactivation and the frequency of Ly6G+ and Ly6C+ cells in the PLF following reactivation (G: p = 0.321; H: p = 0.0025; I: p = 0.0274). **(J)** Flow plots showing gating of CD11b and Ly6G vs Ly6C of Control (top) vs Ablated (bottom). Adjunct histograms on flow plots show the MFI of each marker. D-I: Control n = 11, Ablated n = 14. *Data presented as mean ± SEM. Statistical significance determined by unpaired t-tests and outliers were identified using the ROUT method (Q = 1%) and excluded*.

We could confirm successful reactivation by evaluating c-Fos expression of the TRAPed neuronal population (Fig. 3B). Importantly, RTX treatment had no effect on the number of mCherry+ or c-Fos+mCherry+ TRAPed neurons, indicating that neuronal ensembles were successfully reactivated across groups (Fig. 3C, D).

To assess how TRPV1+ neuron ablation affected the nature of the immune response induced by the neuronal reactivation, we used flow cytometry. We analyzed the composition of immune cells recruited following the reactivation of the pIC ensembles in both groups (Ablated and Control) and performed PCA on immune cell frequencies in the PLF. This revealed a significant separation between groups following reactivation of pIC neurons, indicating a global shift in immune state due to TRPV1+ sensory input (p = 0.0012) (Fig. 3E). Groups separated primarily along PC2 (26.8% of variance), driven by opposing loadings of CD19+MHCII+ and Ly6C+ cells. PC1 (48.1% of variance) reflected within-group differences in myeloid abundance, with F4/80+, CD11c+, and CD11b+ cells as the main contributors.

tSNE embedding of all CD45+ PLF cells pointed to reduced myeloid (CD11b+) frequencies and a relative enrichment of naïve B cells (CD19+) as the cellular basis of this shift (Fig. 3F, Fig. S5). Ablated mice showed markedly fewer reactivation-induced CD11b+ myeloid cells and lower Ly6G+ and Ly6C+ frequencies (Fig. 3G-I). Notably, these effects were male-specific, females showed no difference between Ablated and Control groups at reactivation (Fig. S6). This suggests that females engage distinct encoding mechanisms or compensatory circuits, consistent with established sex differences in pain processing ^37,38^. For that reason, all subsequent experiments were performed on male mice. Taken together, our results dissociate the acute immune response from its neural encoding, demonstrating that TRPV1⁺ neurons are required for the pIC to encode an immune response that can later be reactivated.

### Nociception is required for encoding of inflammatory responses

We next asked whether this requirement reflects a broader role for nociceptive signaling and pain perception, rather than a specific dependence on TRPV1⁺ neurons. To test this, we induced analgesia during the inflammatory episode using a non-inflammatory dose of acetaminophen (Fig. 4A). Acetaminophen mediates analgesia predominantly through central mechanisms, acting within the spinal cord and brain rather than at peripheral nociceptive terminals, including engagement of cannabinoid and serotonergic pathways and suppression of prostaglandin synthesis^39,40^. We confirmed the analgesic efficacy of acetaminophen using the Von Frey test (p = 0.0108) (Fig. 4B).

**Fig. 4.**
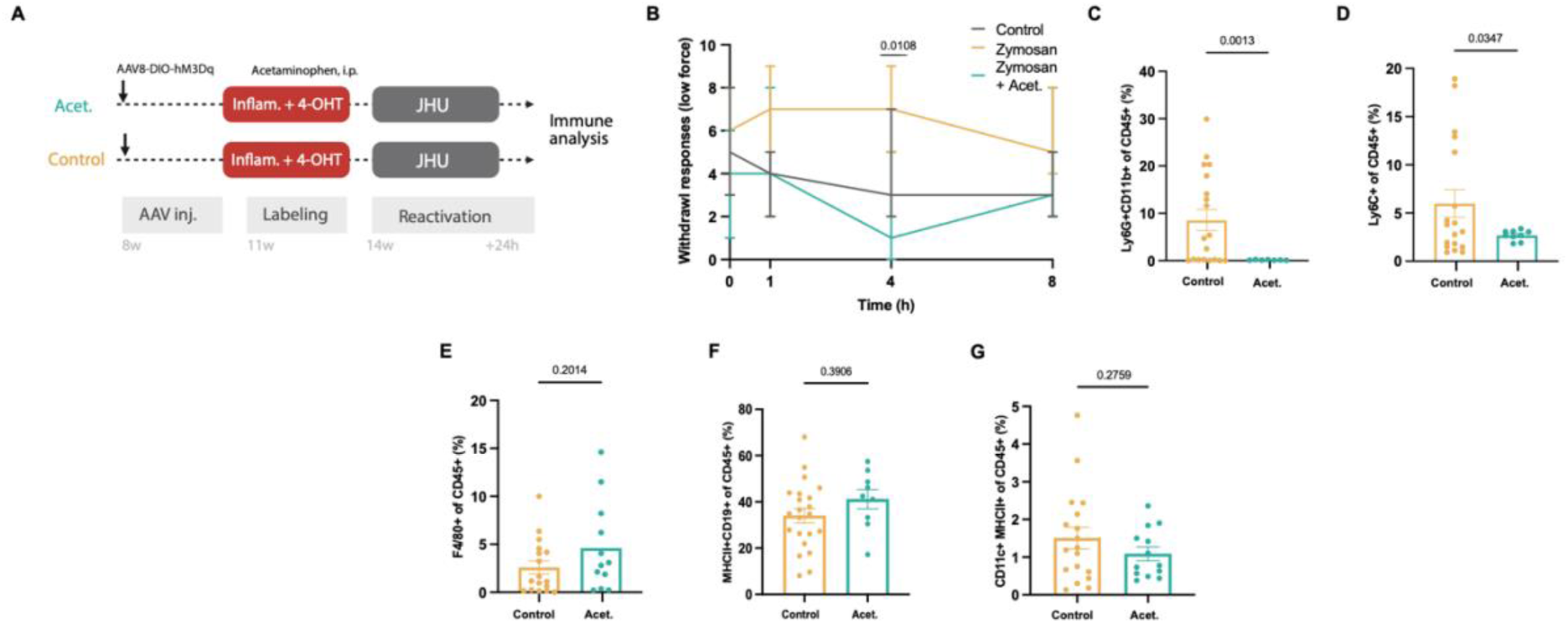
Pharmacological analgesia during inflammation prevents the neural encoding of immune responses. The neural representation of inflammation in the pIC is contingent upon nociceptive signaling; alleviating pain during the initial challenge decouples the immune response from its cortical signature, preventing subsequent immune recall. **(A)** Experimental paradigm for pharmacological pain modulation. Schematic illustrating the administration of the analgesic acetaminophen (300 mg/kg, i.p.) or vehicle during ZIP. Neuronal ensembles in the pIC were TRAPed during this "attenuated-pain" window and reactivated 3 weeks later with JHU. **(B)** Validation of analgesic efficacy. Assessment of pain-related behavior using Von Frey filaments at 1, 4, and 8 hours post-inflammation. Acetaminophen-treated mice exhibited significantly lower low-threshold withdraw responses compared to vehicle-treated inflamed mice (p = 0.0108). Control n =6, Zymosan n = 6, Acet n = 6. **(C-G)** Bar plots of the frequency of cell types in the PLF following reactivation (C: p = 0.0013; D: p = 0.0347). C-H: Control n = 18, Acet n = 8. *Data presented as mean ± SEM. Statistical significance determined by unpaired t-test: B, E-G. Outliers were identified using the ROUT method (Q = 1%) and excluded*.

Consistent with TRPV1⁺ neuron ablation, acetaminophen administration did not alter the acute inflammatory response in the peritoneum (Fig. S7). To capture the immune response during the analgesic time-window, we performed flow cytometric analysis of the PLF 4 hours after zymosan injection. This revealed no significant differences in Ly6G+ or Ly6C+ recruitment between vehicle and acetaminophen-treated mice (Fig. S7), indicating that the peripheral immune response remained intact despite reduced nociceptive input.

To test the effect of analgesic treatment during inflammation on the encoding and recall of inflammation, we again combined activity-dependent neuronal labeling with chemogenetic reactivation in the pIC of mice treated with or without acetaminophen (Fig. 4A). After recovery (3-4 weeks) we administered JHU to activate the TRAPed neurons and assessed the immune response in the PLF 4 hours later by flow cytometry, to match the TRAPing time window.

Subsequent chemogenetic reactivation of inflammation-associated pIC ensembles revealed that analgesia during the initial inflammatory episode attenuated the retrieval capacity of an immune response. Flow cytometric analysis showed a significant reduction in the frequency of Ly6G+ and Ly6C+ cells compared to controls (p = 0.0013 and 0.0347, respectively) (Fig. 4C, D). No changes were observed in F4/80+, CD19+, or CD11c+ populations (Fig. 4E-G). Thus, in the control group not treated with acetaminophen, pIC activation of TRAPed neurons induced immune infiltration into the peritoneum; in mice treated with acetaminophen during the original inflammation, pIC activation did not induce this immune response. These results further demonstrate that pain signaling during inflammation is required for recall of the immune response. It also suggests that analgesic treatment during inflammatory events can alter how the brain represents these immune states and potentially replays them.

### TRPV1⁺ neuron activation reshapes immune recall

If reducing nociceptive signaling attenuates the brain’s immune representation, augmenting it should have the opposite effect by strengthening the encoding of inflammation in the brain. This prediction is consistent with our hypothesis that nociceptive input serves as a salience signal during inflammation. Pain may mark the event as biologically significant and gate the formation of a lasting neural representation. Salient stimuli are well-established drivers of stronger memory encoding across biological contexts^41–43^. If nociception serves this function, then the intensity of the pain signal should actively scale the strength of the encoded immune representation. To test this directly, we transiently activated TRPV1⁺ neurons during the inflammatory episode and asked whether augmented nociceptive input reshapes subsequent immune recall.

Mice received a low dose of RTX (3 µg/kg i.p.) ten minutes before zymosan administration to activate TRPV1⁺ neurons (Fig. S8A). Grimace scores were elevated 15 minutes after zymosan injection in RTX-treated animals (“Activated”), confirming augmented nociceptive signaling during the inflammatory episode (Fig. 5B). This manipulation had minimal effects on the acute inflammatory response, as immune profiling 24 hours after ZIP revealed comparable cellular composition in the PLF between activated and control mice, with only a small but significant decrease in Ly6C+ frequency (p = 0.008) (Fig. S8B-D).

**Fig. 5.**
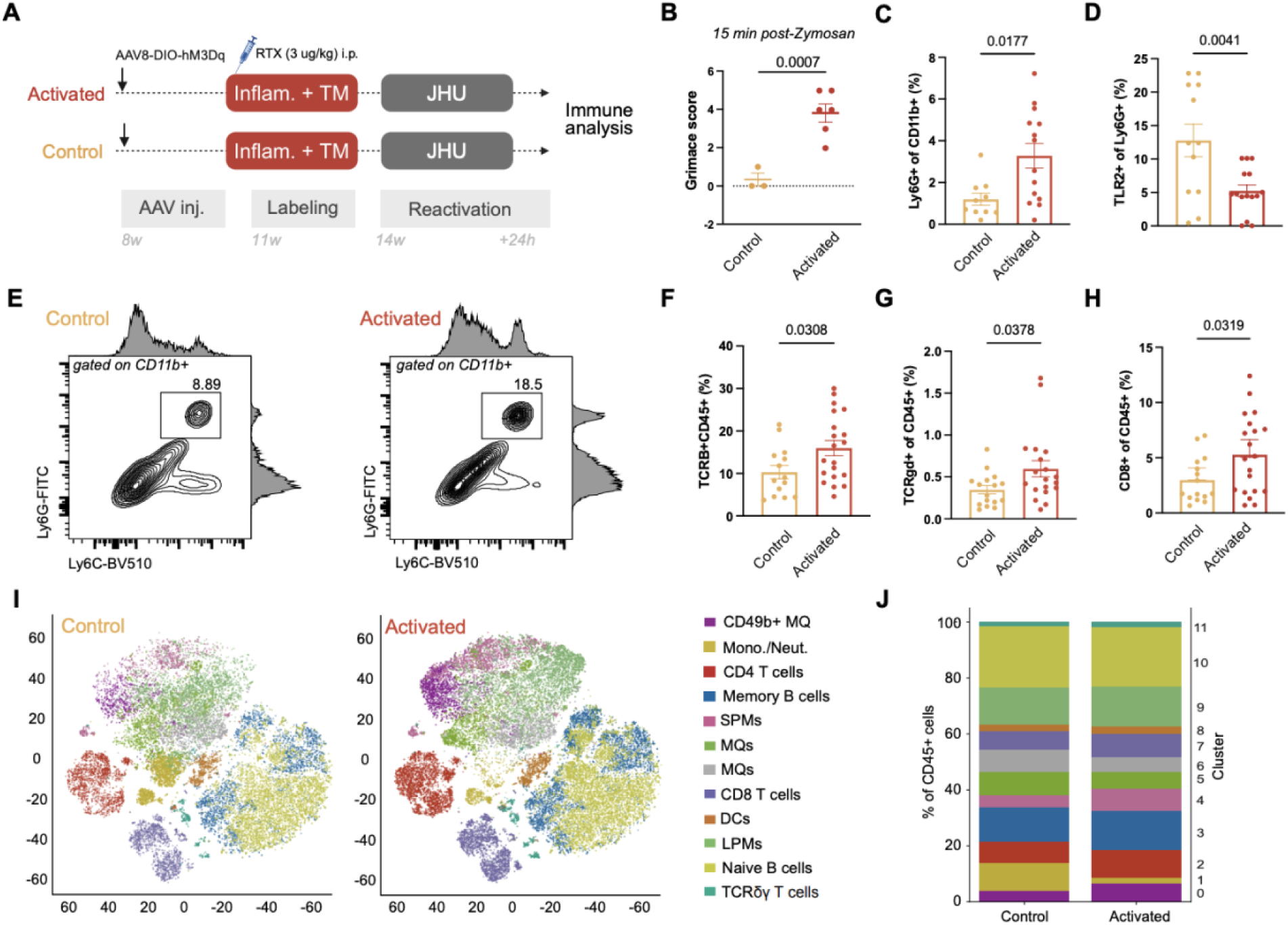
Transient nociceptor activation amplifies the magnitude and shifts the quality of immune recall. The intensity of sensory input during an immune challenge acts as a gain-control mechanism for the insular cortex; heightened nociceptive signaling during initial exposure programs a more potent and qualitatively distinct immune recall that favors lymphoid recruitment upon reactivation. **(A)** Gain-of-function experimental design. Mice received a low dose of RTX (3ug/kg i.p.) 15 minutes prior to ZIP. Controls received vehicle + ZIP. pIC ensembles were TRAPed during this heightened pain state and reactivated 3 weeks later. **(B)** Pain phenotype assessed by the Mouse Grimace Scale (MGS) 15 minutes post-zymosan. RTX-activated mice scored higher than controls (p = 0.0007). Control n = 3, Activated n = 6. **(C)** Frequency of Ly6G+ cells of CD11b+ in the PLF (p = 0.0177). **(D)** Frequency of TLR2+ cells of Ly6G+ in the PLF (p = 0.0041). **(E)** Representative flow cytometry plots showing gating of Ly6G+ cells in Control vs Ablated (Ly6G vs Ly6C), gated on CD11b+. Adjunct histograms on flow plots show the MFI of each marker. **(F)** Frequency of TCRß+ cells of CD45+ (p = 0.0308). **(G)** Frequency of TCRγδ + cells of CD45+ (p = 0. 0378). **(H)** Frequency of CD8 + cells of CD45+ (p = 0. 0319). **(I)** tSNE embedding of CD45+ cells from the PLF in Control and Activated groups. **(J)** Stack bar plots of the relative frequency of each tSNE cluster of out CD45+ cells. Control n = 16, Activated n = 19. *Data presented as mean ± SEM. Statistical significance determined by unpaired t-test. Outliers were identified using the ROUT method (Q = 1%) and excluded*.

Then, as in previous experiments, we combined activity-dependent labeling with chemogenetic reactivation of pIC neurons that were TRAPed during inflammation at baseline or with the increased pain stimulus (Fig. 5A). Indeed, upon chemogenetic reactivation, RTX-activated mice mounted a qualitatively different immune response from controls. As expected, for some immune parameters, such as Ly6G+ cell frequency, we observed an increase in the Activated group compared to controls (p = 0.0177) (Fig. 5C). However, it was not general activation, since, for example, TLR2 expression on these cells was reduced (p = 0.0041) (Fig. 5D–E). We also saw that reactivation of the immune engrams acquired in mice that received increased nociceptive signaling (Activated) led to increased abundance of CD8+TCRß+ and γδ T cells (TCRß+: p = 0.0459; γδ: p = 0.0378; CD8+: p = 0.0319) (Fig. 5F-H). This means that reactivating pIC neurons that were originally active during a more painful inflammation is sufficient to reconstitute a distinct and potentially stronger peripheral immune phenotype. Interestingly, we observed that in the Activated mice, the CD62L+ pool was expanded across CD4+, CD8+, and γδ T cell subsets (p = 0.0345, 0.0216, and 0.0083, respectively) (Fig. S8M-O, left). In CD8+ T cells, this coincided with reductions in both the CD62L+CD44+ and CD62L−CD44+ pools (p = 0.0189 and 0.0464, respectively) (Fig. S8N, middle and right). In CD4+ T cells, the CD62L+CD44+ pool trended higher in activated animals (p = 0.0644) (Fig. S8), middle). Classically, CD62L marks naïve and central memory T cells that home to lymph nodes, while CD44 marks antigen-experienced/activated cells. These results were surprising given that our model is acute and myeloid-driven. No other experimental manipulation in this study produced such extensive effects on the adaptive immune compartment. Thus, this data further suggests that, when pain signaling was increased during the inflammation, not only did the brain’s representation of the immune response become potentially more intense, as evidenced by greater myeloid recruitment, but it also shifted the representation toward a more naïve phenotype of the T cell compartment.

While uncovering the functional meaning of these findings is beyond the scope of this study, our data demonstrates that pIC neurons encode not only whether inflammation occurred, but how intensely it was perceived, with nociceptive input acting as a gain control that sets the balance of subsequent immune recall.

## DISCUSSION

Our findings demonstrate that nociceptive input is required for the brain to encode a retrievable representation of inflammation. We show that peripheral inflammation leads to reorganization of brain network architecture in both humans and mice, shifting from specialized, segregated networks towards a highly integrated brain-wide state. This reorganization depends on intact nociceptive signaling.

Within this reorganization, subcortical circuits remained active in nociceptor-ablated animals, but cortical engagement was selectively lost. This suggests that the brain can mount a peripheral immune response without cortical connectivity, as evidenced by RTX-treated animals maintaining their peripheral immunity. But without cortical engagement, the inflammation was not adequately represented in the brain and could not be retrieved by the reactivation of the immune ensembles in the insular cortex. This is in line with cortical networks being required to transform a peripheral signal into a retrievable state ^44,45^. Thus, we suggest that nociception transforms inflammation from a subcortical reflex into a cortically encoded and retrievable experience, “immune engram”^46^. While we focused on the posterior insular cortex, a key hub for interoceptive, nociceptive, and salience processing^36,47,48,49^ and an established site of immune engram formation^11^, the brain-wide network reorganization we observed suggests that immune encoding may not be restricted to this region alone. Independent evidence similarly implicates TRPV1⁺ sensory neurons in gating insular cortex plasticity following peripheral immune challenge, through a distinct, inhibitory-microcircuit-based mechanism (Z. H. Gursky, E. Favuzzi et al., personal communication).

Memory systems are inherently selective. What determines whether an inflammatory experience is encoded at all? The brain cannot store all physiological fluctuations but should somehow mark biologically relevant events for retrieval. In general, saliency is a critical factor for determining memory formation^50^. Pain is among the strongest salience signals ^8,35,36^, which would explain why nociceptive input is required. Thus, we suggest that pain flags the inflammatory event as significant and recruits the cortical circuits needed to encode it. Indeed, our data are consistent with this interpretation and can explain why the absence of pain, despite an intact peripheral immune response, is sufficient to prevent immune recall.

This nociceptive mechanism is not restricted to TRPV1⁺ neurons. Pharmacological analgesia with acetaminophen, which acts centrally, produced the same failure to encode a retrievable immune representation. The reciprocal is also true; transiently amplifying nociceptive input during inflammation with low-dose RTX strengthened recall and qualitatively reshaped it, shifting the recalled immune response toward greater myeloid engagement and a redistribution in T cell pools. Nociceptive input thus acts as a gain control, setting not only whether an immune representation forms, but what that representation encodes. It is important to note, however, that nociceptive signaling is itself highly complex, and our findings address only one dimension of it. Inflammation sensitizes afferent populations beyond TRPV1⁺ nociceptors, including mechanoreceptors, thermoreceptors, pruriceptors, and silent nociceptors. Whether these parallel sensory inputs also contribute to immune engram formation is not addressed by our experiments. Dissecting which specific aspects of nociception contribute to which features of immune representation remains an important direction for future investigation.

Our findings carry significant clinical implications. Patients with chronic pain show measurable immune dysregulation ^21,51^, though this can arise through multiple mechanisms, not all nociceptive. Particularly compelling is evidence that pain-mediated neural signals can induce disease relapse, demonstrated experimentally in a murine model of multiple sclerosis^52^ and suggested in other autoimmune conditions^53^, pointing to a broader principle whereby the brain’s encoding of painful immune experiences shapes future inflammatory responses. It also suggests that analgesic treatment during acute infection and inflammation may have clinical implications that have not been examined.

Thus, our findings reveal a previously unrecognized function of nociceptive signaling in neuroimmune regulation. Beyond alerting the organism to tissue damage and driving protective behavior, nociceptive input during inflammation shapes how the brain encodes immune events, determining whether, and in what form, an immune response can later be recalled. The implications extend to the management of pain in inflammatory and infectious conditions, where the long-term neuroimmune consequences of analgesia remain largely unknown.

## Acknowledgments

Portions of the manuscript text and analysis code were drafted or edited with the assistance of AI language models (Claude, ChatGPT). All AI-generated content was reviewed, verified, and revised by the authors, who take full responsibility for the accuracy and integrity of the work.

## Author contributions

Conceptualization: MS, AR, IC

Methodology: MS

Investigation: MS

Visualization: MS

Funding acquisition: AR

Project administration: MS

Supervision: AR

Writing – original draft: MS, AR

Writing – review & editing: MS, AR, IC, NS

## Data, code, and materials availability

“All data are available in the main text or the supplementary materials.”

## Supplementary Materials

Materials and Methods Figs. S1 to S8

References (*1*–*49*)

## References and Notes

1. Raja, S. N., et al. The revised International Association for the Study of Pain definition of pain: concepts, challenges, and compromises. Pain 161, (2020).

2. Baral, P., Udit, S. & Chiu, I. M. Pain and immunity: implications for host defence. Nat. Rev. Immunol. 19, 433–447 (2019).

3. Deng, L., Gillis, J. E., Chiu, I. M. & Kaplan, D. H. Sensory neurons: An integrated component of innate immunity. Immunity 57, 815–831 (2024).

4. Pinho-Ribeiro, F. A., Verri Jr., W. A. & Chiu, I. M. Nociceptor Sensory Neuron&#x2013;Immune Interactions in Pain and Inflammation. Trends Immunol. 38, 5–19 (2017).

5. Coghill, R. C. The Distributed Nociceptive System: A Framework for Understanding Pain. Trends Neurosci. 43, 780–794 (2020).

6. Martucci, K. T. & Mackey, S. C. Neuroimaging of Pain: Human Evidence and Clinical Relevance of Central Nervous System Processes and Modulation. Anesthesiology 128, (2018).

7. Kucyi, A. & Davis, K. D. The dynamic pain connectome. Trends Neurosci. 38, 86– 95 (2015).

8. Hemington, K. S., Wu, Q., Kucyi, A., Inman, R. D. & Davis, K. D. Abnormal cross-network functional connectivity in chronic pain and its association with clinical symptoms. Brain Struct. Funct. 221, 4203–4219 (2016).

9. Wager, T. D., et al. An fMRI-Based Neurologic Signature of Physical Pain. New England Journal of Medicine 368, 1388–1397 (2013).

10. Woolf, C. J. & Ma, Q. Nociceptors&#x2014;Noxious Stimulus Detectors. Neuron 55, 353–364 (2007).

11. Koren, T., et al. Insular cortex neurons encode and retrieve specific immune responses. Cell 184, 5902–5915.e17 (2021).

12. Adaikkan, C. & Rosenblum, K. A molecular mechanism underlying gustatory memory trace for an association in the insular cortex. Elife 4, e07582 (2015).

13. Ryan, T. J., Roy, D. S., Pignatelli, M., Arons, A. & Tonegawa, S. Engram cells retain memory under retrograde amnesia. Science (1979). 348, 1007–1013 (2015).

14. Labrenz, F., et al. Alterations in functional connectivity of resting state networks during experimental endotoxemia - An exploratory study in healthy men. Brain Behav. Immun. 54, 17–26 (2016).

15. Lekander, M., et al. Intrinsic functional connectivity of insular cortex and symptoms of sickness during acute experimental inflammation. Brain Behav. Immun. 56, 34– 41 (2016).

16. Färber, N., Manuel, J., May, M., Foadi, N. & Beissner, F. The Central Inflammatory Network: A Hypothalamic fMRI Study of Experimental Endotoxemia in Humans. Neuroimmunomodulation 29, 231–247 (2021).

17. Schaefer, A., et al. Local-Global Parcellation of the Human Cerebral Cortex from Intrinsic Functional Connectivity MRI. Cerebral Cortex 28, 3095–3114 (2018).

18. Tian, Y., Margulies, D. S., Breakspear, M. & Zalesky, A. Topographic organization of the human subcortex unveiled with functional connectivity gradients. Nat. Neurosci. 23, 1421–1432 (2020).

19. Yeo, B. T. T., et al. The organization of the human cerebral cortex estimated by intrinsic functional connectivity. J. Neurophysiol. 106, 1125–1165 (2011).

20. Sporns, O. & Betzel, R. F. Modular Brain Networks. Annu. Rev. Psychol. 67, 613– 640 (2016).

21. Dantzer, R., O’Connor, J. C., Freund, G. G., Johnson, R. W. & Kelley, K. W. From inflammation to sickness and depression: when the immune activation promotes negative affect. Nat Rev Neurosci 9, 46–56 (2008).

22. Struller, F., et al. Peritoneal innervation: embryology and functional anatomy. 2, 153–161 (2017).

23. Peeters, P. J., et al. Molecular profiling of murine sensory neurons in the nodose and dorsal root ganglia labeled from the peritoneal cavity. Physiol. Genomics 24, 252– 263 (2006).

24. Cavanaugh, D. J., et al. Restriction of Transient Receptor Potential Vanilloid-1 to the Peptidergic Subset of Primary Afferent Neurons Follows Its Developmental Downregulation in Nonpeptidergic Neurons. The Journal of Neuroscience 31, 10119 (2011).

25. Caterina, M. J., et al. Impaired Nociception and Pain Sensation in Mice Lacking the Capsaicin Receptor. Science (1979). 288, 306–313 (2000).

26. Maximiano, T. K. E., Carneiro, J. A., Fattori, V. & Verri, W. A. TRPV1: Receptor structure, activation, modulation and role in neuro-immune interactions and pain. Cell Calcium 119, 102870 (2024).

27. Mishra, S. K., Tisel, S. M., Orestes, P., Bhangoo, S. K. & Hoon, M. A. TRPV1-lineage neurons are required for thermal sensation. EMBO J. 30, 582–593 (2011).

28. Palazzo, E., et al. Transient receptor potential vanilloid type 1 and pain development. Curr. Opin. Pharmacol. 12, 9–17 (2012).

29. Guenthner, C. J., Miyamichi, K., Yang, H. H., Heller, H. C. & Luo, L. Permanent Genetic Access to Transiently Active Neurons via TRAP: Targeted Recombination in Active Populations. Neuron 78, 773–784 (2013).

30. Mitchell, Kendall et al. Ablation of Rat TRPV1-Expressing Adelta/C-Fibers with Resiniferatoxin: Analysis of Withdrawal Behaviors, Recovery of Function and Molecular Correlates. Mol. Pain 6, 1744-8069-6–94 (2010).

31. Dale, J., et al. Scaling Up Cortical Control Inhibits Pain. Cell Rep. 23, 1301–1313 (2018).

32. Urien, L. & Wang, J. Top-Down Cortical Control of Acute and Chronic Pain. Biopsychosocial Science and Medicine 81, (2019).

33. Liang, D. & Labrakakis, C. Multiple Posterior Insula Projections to the Brainstem Descending Pain Modulatory System. Int. J. Mol. Sci. 25, 9185 (2024).

34. Buzsáki, G. Neural Syntax: Cell Assemblies, Synapsembles, and Readers. Neuron 68, 362–385 (2010).

35. Borsook, D., et al. The Insula: A ‘Hub of Activity’ in Migraine. Neuroscientist 22, 632–652 (2016).

36. Legrain, V., Iannetti, G. D., Plaghki, L. & Mouraux, A. The pain matrix reloaded: A salience detection system for the body. Prog. Neurobiol. 93, 111–124 (2011).

37. Mogil, J. S. Sex differences in pain and pain inhibition: multiple explanations of a controversial phenomenon. Nat. Rev. Neurosci. 13, 859–866 (2012).

38. Sorge, R. E., et al. Different immune cells mediate mechanical pain hypersensitivity in male and female mice. Nat. Neurosci. 18, 1081–1083 (2015).

39. Dani, C., et al. Intravenous paracetamol in comparison with ibuprofen for the treatment of patent ductus arteriosus in preterm infants: a randomized controlled trial. Eur. J. Pediatr. 180, 807–816 (2021).

40. Ottani, A., Leone, S., Sandrini, M., Ferrari, A. & Bertolini, A. The analgesic activity of paracetamol is prevented by the blockade of cannabinoid CB1 receptors. Eur. J. Pharmacol. 531, 280–281 (2006).

41. McGaugh, J. L. Memory--a Century of Consolidation. Science (1979). 287, 248–251 (2000).

42. LaBar, K. S. & Cabeza, R. Cognitive neuroscience of emotional memory. Nat. Rev. Neurosci. 7, 54–64 (2006).

43. Dunsmoor, J. E., Murty, V. P., Davachi, L. & Phelps, E. A. Emotional learning selectively and retroactively strengthens memories for related events. Nature 520, 345–348 (2015).

44. Damasio, A. & Carvalho, G. B. The nature of feelings: evolutionary and neurobiological origins. Nat. Rev. Neurosci. 14, 143–152 (2013).

45. Dehaene, S. & Changeux, J.-P. Experimental and Theoretical Approaches to Conscious Processing. Neuron 70, 200–227 (2011).

46. Koren, T. & Rolls, A. Immunoception: Defining brain-regulated immunity. Neuron 110, 3425–3428 (2022).

47. McBenedict, B., et al. The Role of the Insula in Chronic Pain and Associated Structural Changes: An Integrative Review. Cureus 16, e58511 (2024).

48. Livneh, Y., et al. Homeostatic circuits selectively gate food cue responses in insular cortex. Nature 546, 611–616 (2017).

49. Livneh, Y. & Andermann, M. L. Cellular activity in insular cortex across seconds to hours: Sensations and predictions of bodily states. Neuron 109, 3576–3593 (2021).

50. McGaugh, J. L. The amygdala modulates the consolidation of memories of emotionally arousing experiences. Annu. Rev. Neurosci. 27, 1–28 (2004).

51. Ji, R.-R., Xu, Z.-Z. & Gao, Y.-J. Emerging targets in neuroinflammation-driven chronic pain. Nat. Rev. Drug Discov. 13, 533–548 (2014).

52. Arima, Y., et al. A pain-mediated neural signal induces relapse in murine autoimmune encephalomyelitis, a multiple sclerosis model. Elife 4, e08733 (2015).

53. Totsch, Stacie K & Sorge, Robert E. Immune system involvement in specific pain conditions. Mol. Pain 13, 1744806917724559 (2017).

