## Supplemental for "Pain Matters: Nociceptive signaling drives neural encoding of inflammatory responses"

Materials and Methods

### Experimental model and subject details

#### fMRI participants, experimental materials and procedure

For the fMRI analysis, we analyzed a previously published dataset [1], comprising 7 healthy adult participants. The study was approved by the Ethics Committee of the Medical Faculty of the University of Duisburg-Essen and all participants provided written informed consent prior to enrollment. Participants received an intravenous bolus of bacterial endotoxin (lipopolysaccharide, LPS; 1.0 ng/kg body weight) to induce a controlled, transient inflammatory response. Resting-state fMRI data were acquired at four time points relative to LPS administration: Baseline (pre-injection), Onset, Acute, and Peak inflammatory phase. Each scan lasted 20 minutes; participants were instructed to remain still with eyes closed and to refrain from falling asleep. The inter-scan interval was approximately 10-20 minutes.

#### Mice

All experiments were performed in accordance with the National Institutes of Health Guide for the Care and Use of Laboratory Animals. All procedures and protocols were approved by the Technion Administrative Panel of Laboratory Animal Care and the Tel Aviv University Administrative Panel of Laboratory Animal Care. Male or female adult (8-12 weeks of age; 20-25 g) FosTRAP2 (Jackson Laboratory; Fostm2.1(icre/ERT2)Luo/J; stock #030323) transgenic mice were used for the InsCtx activity-dependent labeling experiments; Ai14D mice (Jackson Laboratory; B6;129S6-Gt(ROSA)26Sortm14(CAG-tdTomato)Hze/J; Stock #007908) were crossed with FosTRAP2 for whole-brain activity-dependent labeling analysis; and C57BL/6J mice (Jackson Laboratory; C57BL/6J; stock #000664) were used for the characterization of Zymosan-induced peritonitis. Mice were maintained under specific-pathogen-free (SPF) conditions on a 12 h light: 12 h dark cycle (lights on at 07:00) and housed in groups of 2-5 mice per cage, with food and water ad libitum.

### Methods

### fMRI Data Acquisition

Secondary analysis of a published dataset, for full acquisition details, see Färber et al. (2021). Briefly, fMRI data were acquired on a Siemens 3T MAGNETOM Skyra scanner. Functional images were acquired using a gradient-echo EPI sequence with the following parameters: TR = 1,180 ms, TE = 32 ms, simultaneous multi-slice factor = 6; partial Fourier = 7/8; 2 mm isotropic resolution. A T1-weighted structural scan was also acquired for each participant for registration purposes (TR = 2,400 ms; TE = 2.13 ms; TI = 1,000 ms; in-plane acceleration factor = 2; and isotropic resolution: 1 mm).

### fMRI Preprocessing

Functional and structural MRI data were processed using a specialized high-fidelity pipeline (jpreprocessing v2.4) optimized for subcortical and brainstem sensitivity. To minimize spatial blurring and resampling artifacts, susceptibility-induced distortion correction via FSL v5.0.9 topup and retrospective motion correction via MCFLIRT were concatenated into a single-step rigid transformation. Functional data were high-pass filtered at 0.005 Hz to remove low-frequency drift; notably, no spatial smoothing was applied to preserve the high-resolution signal integrity required for fine-grained anatomical parcellation. Subjects' functional scans were registered to a study-specific T1-weighted template using ANTs Symmetric Normalization (SyN) and subsequently transformed into MNI152 standard space to ensure consistent anatomical alignment across the cohort. Six motion parameters derived from rigid-body realignment (three translations, three rotations) were included as nuisance regressors during time series extraction. To minimize physiological noise contamination, mean white matter and CSF signals were extracted from anatomically defined masks and regressed from the BOLD time series prior to connectivity estimation.

### Connectivity Estimation Procedure

Parcel-wise functional connectivity was estimated using the Schaefer 2018 cortical atlas (100 parcels, 7-network solution) [2] combined with the Tian subcortical atlas (Scale I, 16 regions) [3], yielding 116 regions of interest in total. Notably, this clustering assigns specific regions of the anterior cingulate (ACC) and medial prefrontal cortex (mPFC) to the Control network based on large-scale functional connectivity patterns, despite their classical anatomical association with limbic or salience processing. To supplement this, subcortical nodes from the Tian atlas (Scale I) were assigned to networks based on established functional-anatomical roles: the amygdala, hippocampus, and nucleus accumbens were assigned to the Limbic network; the caudate to the Control network; and the thalamus to the Salience network to capture its role as an interoceptive gateway.

For each subject and each scan session, BOLD time series were extracted from each parcel using a NiftiLabelsMasker (nilearn). Pairwise Pearson correlations were computed between all region pairs to construct a 116 × 116 connectivity matrix per subject per timepoint. Correlation coefficients were Fisher Z-transformed prior to group-level analysis to normalize their distribution. Connectivity matrices were computed per subject before group analysis; group-level comparisons were performed on the Fisher Z values.

Brain State Distance

To track the evolution of brain states throughout the inflammatory response, we represented each subject's functional architecture at each phase as a 116-dimensional connectivity vector[4]. This vector was composed of the mean Fisher-z transformed connectivity for each region of interest (ROI) across its entire connectivity profile. We quantified the shift in brain state by calculating the Euclidean distance between a subject's connectivity vector at a given phase and their own baseline. This within-subject approach effectively controls for individual differences in resting-state architecture. Statistical significance for each phase was assessed using one-sample t-tests against zero, and the temporal progression of the shift was validated by fitting linear slopes to individual subject trajectories.

Functional Segregation: Within/Between-Network Ratio

We quantified functional segregation by calculating the ratio of mean within-network connectivity to mean between-network connectivity for each phase[5]. Connectivity values were based on the Fisher-z transformed correlations between ROI pairs assigned to the seven canonical Yeo networks. A ratio above 1 indicates a modular organization with stronger internal network coupling, while values approaching 1 suggest a loss of network differentiation. Each post-baseline phase was compared to baseline using paired t-tests.

Network-Pair Connectivity

To resolve changes in connectivity between specific systems, we computed the mean Fisher-z transformed connectivity for every pairwise combination of the seven networks, including within-network coupling. These relationships were represented in a 7 x 7 matrix for each subject and phase. For every network pair, we calculated the change from baseline (delta-Z) and identified significant shifts using paired t-tests.

TRPV1 Ablation

Resiniferatoxin (RTX, AdioGen Life Sciences), a potent capsaicin analog, was used to deplete TRPV1-positive nociceptors. Male and female 4-week-old mice were given three escalating doses of RTX (30 µg/kg, 70 µg/kg, 100 µg/kg, diluted in PBS with 1.2% DMSO and 0.06% Tween-80) subcutaneously injected over the flank on three consecutive days, as adapted from established protocols[6]. Control littermates were injected with vehicle solution on the same days (PBS with 1.2% DMSO and 0.06% Tween-80). Mice were used for experiments four weeks after the last injection of RTX. Efficiency of RTX treatment in depleting TRPV1-positive nociceptors was confirmed behaviorally by hot plate assay and by counting the number of TRPV1-positive neurons in the DRGs.

Stereotactic injections

Viral vectors and viral vector plasmids were designed and produced by the Viral Vector Facility (VVF) of the Neuroscience Center Zurich. For virus delivery, mice were anesthetized with a ketamine-xylazine mixture (ketamine 80 mg per kg body weight (mg/kg); xylazine 15-20 mg/kg; Sigma-Aldrich) in sterile saline (0.9% NaCl). After shaving and sterilization of the head, mice were placed on a sterile surgical pad on top of a heating pad. Ophthalmic ointment (Duratears, Alcon) was placed over the eyes to prevent dehydration, and the incision site was sterilized. Upon loss of recoil paw compression, a midline incision was made from above the forehead to the ears line, exposing the skull. After skull exposure, mice were fixed in the stereotactic frame (Stoelting) and carefully drilled through the skull, just above the target injection site. Using a 10 µL syringe with a blunt metal needle (Nanofil), placed in a Stoelting Quintessential Stereotaxic Injector (QSI), we validated that the skull was straight at all plains. We then slowly lowered the syringe into the injection site and left the needle in place for 5 minutes, both before and after the injection. We injected 0.40 µL of virus (either AAV8-hSyn-DIO-hM3D(Gq)-mCherry or AAV8-hSyn-DIO-mCherry) at 0.07 µL per min into the right posterior InsCtx (AP, −0.35 mm; ML, 4.0 mm; DV, −3.83 mm; relative to bregma). Following injection, the incision was closed using a tissue adhesive (Histacryl) and sterilized with Polydine ointment. Mice showing signs of physical distress and pain were excluded from the experiment. Further procedures of the experiments were performed at least 21 days after surgery to ensure the expression of the virus and to avoid analyzing the immediate immune response that could be induced by the injection itself. Stereotactic injection sites were verified using fluorescence microscopy.

Zymosan-induced peritonitis

For peritonitis induction, we used Zymosan A prepared from Saccharomyces cerevisiae cell wall (Sigma-Aldrich). 20 mg of Zymosan were dissolved in 1 mL of sterile HBSS (without $Ca^{+2}$ and $Mg^{+2}$, Biological Industries), during heating in a boiling water bath for 20 min. We sonicated the solution for 20 min, washed twice with HBSS (1200 rpm, 10 min) and resuspended in HBSS to a concentration of 20 mg/ml. The Zymosan solution was aliquoted and kept at −80°C. Upon use, the Zymosan solution was diluted in sterile PBS to a concentration of 20 µg/ml and a volume of 0.25 mL was administered to each mouse via intraperitoneal (i.p.) injections (100 µg of Zymosan per peritoneal cavity). As previously described[7], this dose is sufficient to induce a well-characterized inflammatory response, which is confined to the peritoneal cavity, and resolves within 48 h after induction.

Activity-dependent cell labeling

For ablation or activation experiments, Tamoxifen (TM, Sigma-Aldrich) was dissolved in corn oil (Sigma-Aldrich) to a concentration of 20 mg/ml and kept at 4°C until use (up to 12 h). To enable a Cre-dependent expression of the target gene (either Gq-DREADD or fluorescent reporter) in active neurons, 150 mg/kg of the TM solution was injected i.p. 48 h or 2 h before Zymosan administration (for capturing InsCtx neurons active during a healthy state or peritonitis, respectively).

For pharmacological analgesia experiments, 4-Hydroxytamoxifen (4-OHT, Sigma-Aldrich) was dissolved fresh at 20 mg/ml in ethanol at 37°C. Corn oil (Sigma-Aldrich) was added to give a final concentration of 10 mg/ml 4-OHT, and the ethanol was evaporated by vacuum using centrifugation. A dose of 50 mg/kg of the final 4-OHT solution was injected i.p. 48 h before or right before Zymosan administration. This was used to label neurons only during the active analgesic time-window.

Chemogenetic neuronal manipulation

To activate DREADD-expressing neurons in the InsCtx, all mice were injected i.p. with JHU37160 (JHU) (0.1 mg/kg; Hello Bio) in sterile saline, 3-4 weeks after TM or 4-OHT administration. A single dose of JHU was administered 4-24 h prior to euthanizing the mice, depending on the experiment. For c-Fos evaluation, an additional JHU injection was administered 90 min before euthanasia. Of note, in all experiments, mice in all groups were injected with JHU to control for its potential effects[8].

Tissue preparation and immunofluorescence

Validation of the virus injection site, evaluation of DREADDs' activity and c-Fos expression, were performed by immunofluorescence staining and fluorescence and confocal microscopy. Mice were euthanized, and their brains were extracted and fixed in 4% paraformaldehyde (PFA) for 24 h and cryoprotected in 30% sucrose solution for 72 h. Coronal cryosections of the brains were sliced at 50 μm thickness. These were then transferred to cryoprotectant and kept at -20C until used for staining. For immunofluorescence, the mounted tissue sections were incubated in a blocking buffer (0.2% Triton X-100, 0.05% sodium azide, 4% bovine serum albumin in PBS) for 1 h at room temperature. Primary antibodies diluted in blocking buffer were added to the desired concentration and the slides were incubated at 4°C overnight. The following primary antibodies were used for brain sections: chicken anti-mCherry (1:500, Abcam), rabbit anti-cFos (1:2000, Encor). To evaluate c-Fos expression in the brain, mice were sacrificed 90 min after JHU injection and treated as described above. Following overnight incubation with primary antibody, slides were then washed in PBS 3 times for 15 min and further incubated for 1 h at room temperature with a fluorescent-conjugated secondary antibody. The following secondary antibodies were used: Cy5 goat anti-rabbit IgG (1:1000, Jackson ImmunoResearch laboratories); Alexa Fluor-568 goat antichicken (1:1000, Invitrogen). Slides were next incubated with 1ug/mL DAPI in PBS for 5 min at room temperature then washed in PBS 3 times for 15 min. These were then mounted on super-frost slides (Thermo scientific) with mounting medium (SouthernBiotech) and covered with a glass coverslip (Bar-Naor). All images were taken at 10 × or 20 × magnification using the Leice SP8 confocal microscope.

Whole brain clearing

Whole brain clearing was performed as previously described, with some adjustments[9]. Briefly, after 4% PFA fixation, brains were dehydrated with increasing concentrations of methanol in B1n buffer[10], and incubated overnight in 2:1 DCM:MeOH. The following day, the brains were washed in 2xDCM, 2xMeOH and then incubated with 5% H2O2 in methanol (MeOH) at 4°C overnight. The following day, they were washed 2xMeOH, then rehydrated in decreasing concentrations of MeOH/B1nSamples, and were then incubated in permeabilization buffer for 2-3 days in a 37C water bath. Primary antibody was added in PTwH (rabbit anti-RFP, 1:300, Rockland ) for 11 days in 37C then washed 4-5 times overnight in PTwH. Secondary antibody was added in PTwH (donkey anti-rabbit AF647, 1:300, Invitrogen) for 8 days in 37C then washed 4-5 times and overnight in PTwH. Finally, brains were dehydrated in increasing concentrations of MeOH in DDW, 2xMeOH 100%, 2:1 DCM:MeOH solution at room temperature for 3 hours, and then in 100% DCM for 15 min twice, while shaking. Tissues were stored in refractive index (RI) matching solution (RI = 1.56), Dibenzyl ether (DBE, Sigma-Aldrich) until imaging.

Light-sheet imaging

Whole-brain cleared samples were imaged while submerged in DBE in the 3iCTLS light sheet in the large chamber, using 561nm and 640nm illumination, 1x objective, and 4µm z-slices, at a resolution of 1.97 µm (xy) and 4.0 µm (z).

Whole-brain analysis

Raw whole-brain volumes were processed using TRAILMAP and DeepTraCE, deep-learning based pipelines for analysis of whole-brain images. Cell bodies were segmented in the raw coordinate space using TRAILMAP[11]. Then, volumes were scaled to 10 µm isotropic and 8-bit format and registered to the Allen Mouse Brain Common Coordinate Framework (CCFv3) at 25 μm resolution using the brainreg pipeline[12]. Precise 3D rigid-body rotations were applied to correct for sample-to-atlas orientation bias, with the exact transformation matrices co-applied to the TRAILMAP binary masks to maintain spatial synchronicity.

In other experiments, whole-brain coronal sections of 40 μm were imaged at 10x and processed DMC-BrainMap, a napari plugin for whole-brain analysis. Preprocessing included padding and resizing of images to ensure consistent dimensions across sections. Channel-specific images were combined to generate RGB composites for visualization and converted into a format compatible with the SharPy tracking framework for registration. Registration was performed using the SHARP-Track workflow. Individual sections were manually aligned to the CCFv3 (25 μm resolution) using anatomical landmarks. Transformations were saved and applied across sections to map image data into atlas space. Following registration, anatomical region assignments were obtained based on the Allen Brain Atlas ontology, enabling region-level quantification of signal distribution. Quantification of cells per brain calculated as total number of active cells.

Whole-brain co-activation matrices

To characterize the functional organization of the brain during inflammation, we constructed region-by-region co-activation matrices for each experimental group. These matrices were built by computing Pearson correlations of tdTomato+ cell densities between all brain regions across animals.

Hierarchical clustering and module detection

We identified discrete functional modules using agglomerative hierarchical clustering on the correlation-distance matrix with average linkage (UPGMA). The number of clusters was set to k = 5 based on the maxclust criterion, which showed the strongest correspondence with major anatomical systems and was confirmed by stability analysis. Clustering was performed on pooled samples and then groups were separately plotted for visualization.

Cluster stability

We validated the robustness of our 5-cluster partition using a bootstrap stability analysis (100 iterations). Animals were resampled with replacement, and cluster consistency was quantified using the Jaccard similarity index. All five clusters demonstrated high stability, with mean Jaccard indices of approximately 0.90, exceeding the predefined stability threshold of 0.85.

Regional brain system enrichment

We used Fisher's exact tests (one-sided) to determine if specific functional clusters were significantly enriched with regions from particular anatomical parent systems. P-values were FDR-corrected across all tests. In bubble plot visualizations, bubble size represents the region count, while opacity indicates statistical significance ($q < 0.05$). Fiber tracts and Unknown/Other parent categories were excluded from enrichment tests.

Within-cluster connectivity distributions

To assess the global coherence of each module, we calculated the mean intra-cluster Pearson correlation (r) for every animal. Differences in distribution between groups were analyzed using the Mann-Whitney U test.

Flow cytometry

Mice were euthanized, and their peritoneal lavage fluid (PLF) and spleen were collected. Peritoneal cells were collected via peritoneal lavage; 5 mL of cold PBS was carefully injected into the peritoneum, and the cell-containing fluid was collected with a Pasteur pipette into 15 mL tubes. These were centrifuged at 1200 rpm for 5 min. PLF supernatant was extracted and frozen at −20°C for further protein analysis. The recovered cells were incubated for 3 min with 1 mL of Lysis buffer and washed once with PBS. In case of blood-contamination in PLF samples, these samples were excluded from the experiment. Spleen tissue was mechanically dissociated using the back of a syringe in 3 mL of cold PBS in 6 well plates. The suspension was transferred through 40 um mesh strainers into 50 mL tubes and centrifuged at 1200 rpm for 5 min. The recovered cells were incubated for 3 min with 1 mL of Lysis buffer and washed once with PBS. For viability staining, cells (106) were washed once with PBS, resuspended in Zombie Aqua™ dye solution (1:1,000, BioLegend) and incubated in the dark for 15 min at room temperature. For extracellular staining, cells were washed with FACS staining buffer (PBS containing 1% bovine serum albumin, 1 mM EDTA and 0.05% sodium azide) and incubated with antibodies for 30 min at 4 °C. After the incubation of the samples with the antibodies, the samples were washed twice with FACS staining buffer and re-suspended in 250 µl of 1% PFA. Samples were analyzed with a CytoFLEX S or CytoFLEX Mosaic cell analyzer and FlowJo software. For visualization of the immune cell subsets that were analyzed, flow cytometry data was subjected to the unsupervised identification method, FlowSOM. Briefly, we concatenated an equal number of live CD45+ cells from PLF to generate a single FlowSOM tree.

Principal Component Analysis

PCA was run on immune cell frequencies from PLF flow cytometry data. Frequency values were log-transformed and z-scored before analysis (scikit-learn, Python). Group separation was tested by PERMANOVA (adonis2, vegan package, R; 9,999 permutations; Euclidean distance). PERMDISP was run alongside to distinguish true centroid shifts from differences in within-group spread. Confidence ellipses at 68% and 95% were computed assuming bivariate normality.

tSNE analysis

Signal intensities for all markers were normalized using an arcsinh transformation with a cofactor of 150.0 to stabilize variance and mean fluorescence intensity (MFI) values were downsampled to 10,000 cells per sample. The data were then standardized to zero mean and unit variance using the scikit-learn StandardScaler. Data was then embedded in 2-dimensional space using the openTSNE package. Cellular populations were defined via Self-Organizing Map (SOM) clustering using the minisom package. The data were mapped to a 10x10 grid, and the resulting 100 nodes were aggregated into 13 meta-clusters using K-Means clustering.

Von Frey test

Abdominal pain sensitivity was measured using Von Frey filaments as previously described[13]. In brief, mice were randomly divided into 2 groups. One group received an injection of acetaminophen (300 mg/kg) i.p. 15 minutes prior to Zymosan. Another group received an injection of PBS as a control for the analgesia 15 minutes prior to Zymosan. We chose Acetaminophen because it has relatively low anti-inflammatory effects, and it does not block the transition of neuronal inputs to the brain[14]. 10 min later, mice were habituated in metal cubic compartments (13x15 cm2) with a mesh bottom for 15 min. A blinded observer applied Von Frey filaments to the abdomen in ascending order starting at 0.02, 0.04, 0.07, 0.16, 0.4, and 0.6 g forces followed by the same descending order. Each filament was applied three times with maximal duration of 2-3 s, and a 5 s interval between each poke. The number of withdrawal responses (licking, retraction, or abdomen withdrawal) was quantified and summed for each mouse.

Grimace Scoring

Mice were acclimated to custom built aluminum cubic compartments (5x5x5cm) for 30 minutes prior to any injections. Mice were then recorded using GoPro cameras for 30 minutes at baseline, 1, 4, and 8 hours after Zymosan injection. One group was pre-treated with RTX (3 ug/kg) i.p., one with acetaminophen (300 mg/kg) i.p. 15 minutes prior to Zymosan injection. Grimace scoring was then performed by uploading individual videos to PainFace.net[15].

Hot Plate Assay

Thermal nociceptive thresholds were assessed using a hot plate assay maintained at a constant temperature of 57°C. After a 60-minute room habituation period, mice were placed individually onto the heated surface within a transparent plexiglass cylinder. The latency to exhibit nociceptive behaviors, defined as hind-paw licking, flicking, or jumping, was recorded to the nearest second. To prevent thermal tissue injury, a strict 30-second cut-off was enforced, after which any non-responsive mice were immediately removed and assigned a maximum latency score of 30 seconds. The apparatus was cleaned with 30% ethanol between subjects to eliminate olfactory cues.

Illustrations

All figures were created using Illustrator software (Adobe) and BioRender.com

Quantification and statistical analysis

Statistical details for every experiment are provided in the figure legends, where n represents the number of animals per group (unless stated otherwise). Statistical significance was defined as $p < 0.05$. Sample size was chosen based on calibration experiments to detect statistical significance under technical feasibility constrains.

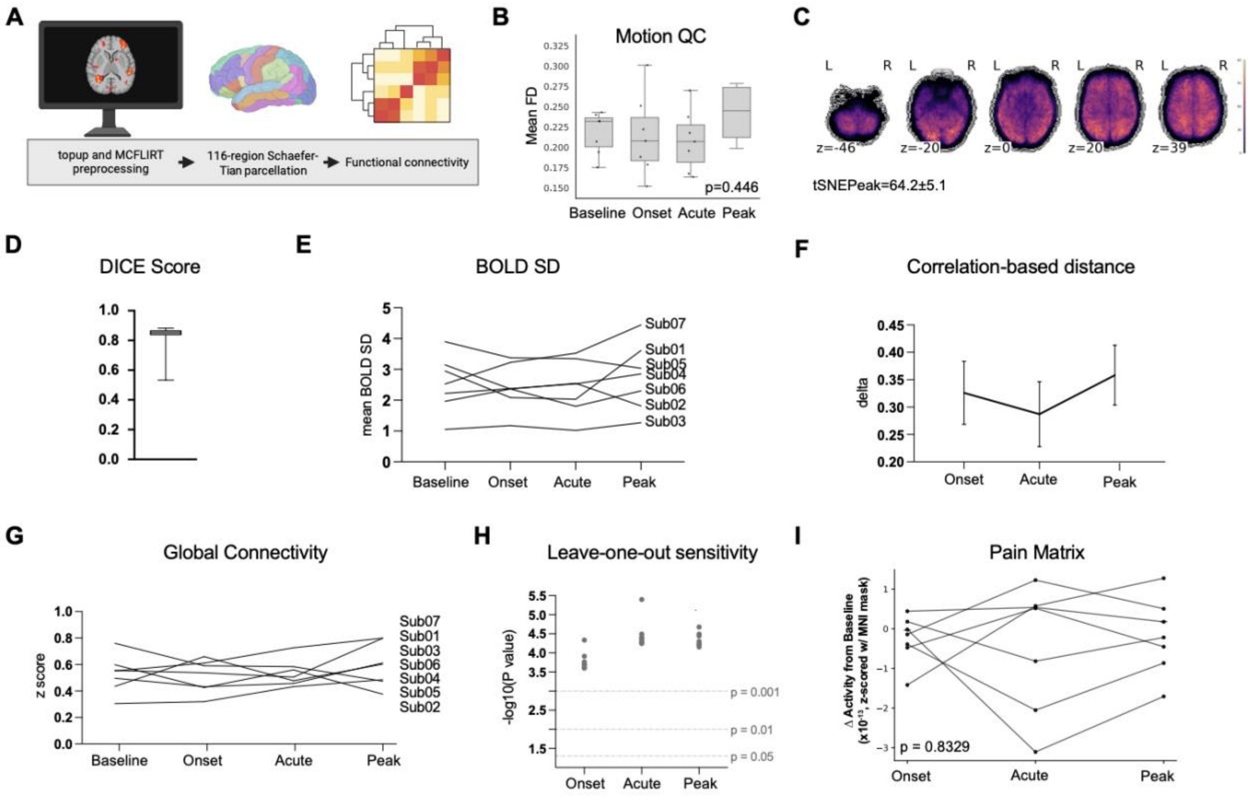


**Fig. S1. fMRI validation**

Technical validation and topological characterization of the human inflammatory brain state. **(A)** Preprocessing pipeline. fMRI data were processed using FSL topup (susceptibility distortion correction) and MCFLIRT (motion correction) as a single-step transformation, parcellated into 116 regions (Schaefer100+Tian16 atlas), and functional connectivity matrices computed from the resulting time series. **(B)** Head motion. Mean framewise displacement (FD) did not differ across phases (p=0.446, one-way repeated-measures ANOVA). **(C)** Temporal SNR. Voxel-wise tSNR map during the Peak phase. Group mean tSNR = 64.2 ± 5.1. **(D)** DICE score. Overlap coefficient between each subject's parcellation mask and the group-average atlas, confirming registration quality (group median DICE > 0.80). **(E)** BOLD signal SD. Standard deviation of the BOLD signal per subject across phases. SD was unchanged at Peak vs. Baseline (paired t-test: t(6) = 0.35, p = 0.600, d = 0.21). **(F)** Correlation-based distance from Baseline, as an independent brain-state metric. Effect sizes were large at Onset (d = 2.14), Acute (d = 1.84), and Peak (d = 2.49), reaching significance at Acute (**p < 0.01) and Peak (***p < 0.001; paired t-tests). **(I)** Global mean connectivity (z-score) per subject across phases. Stable between Peak and Baseline (paired t-test: t(6) = 0.76, p = 0.478, d = 0.29), confirming that the connectivity pattern changes in Fig. 1 do not reflect a global mean shift. **(G)** Leave-one-out sensitivity. Brain-state distance effect (−log10 p) computed from the full cohort (n = 7, black diamonds) and all leave-one-out subsets (n = 6, gray circles) at each phase. **(H)** Activity within the Neurologic Pain Signature (NPS) a whole-brain multivariate pattern trained to discriminate nociceptive from non-painful states (cortical + thalamic MNI mask, Z-scored) plotted as change from baseline for each subject. (repeated measures ANOVA: $F(3,18) = 0.29$, $p = 0.8329$, partial $\eta^2 = 0.046$; n = 7)[16].

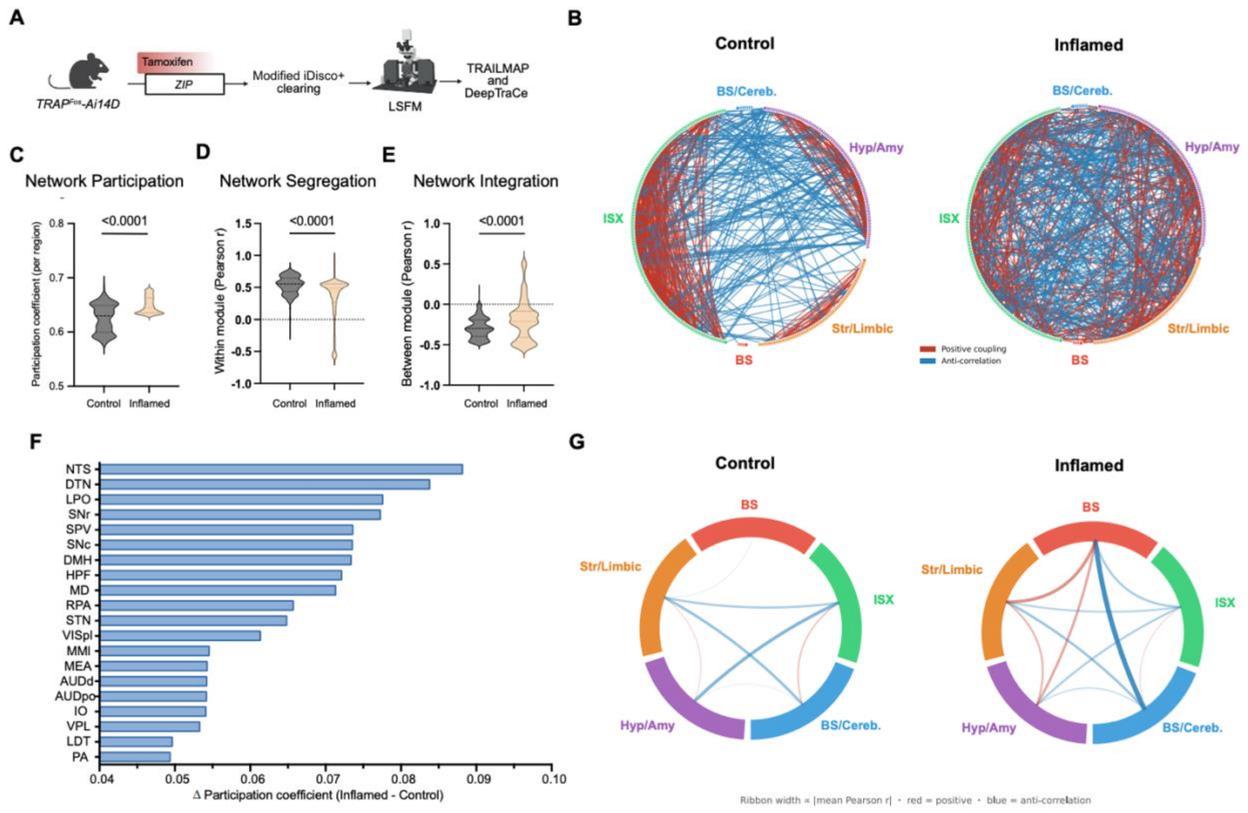


**Fig. S2. Brain-wide network reorganization during sterile peritoneal inflammation.**

**(A)** FosTRAP2-Ai14D mice received tamoxifen during Zymosan-induced peritonitis (Inflamed) or saline injection (Control). Brains were cleared by modified iDisco+, imaged by light sheet fluorescence microscopy (LSFM), and tdTomato+ cells were mapped to Allen Mouse Brain Atlas regions using TRAILMAP and DeepTraCe. **(B)** Circular co-activation networks, each node represents one brain region (n=313), arranged in circular arcs by module: isocortex (ISX, green), hypothalamus/amygdala (Hyp/Amy, purple), striatum/limbic (Str/Limbic, orange), brainstem/cerebellum (BS/Cereb., blue), brainstem (BS, red). Edges connect region pairs with $|r|\geq0.45$ (top ~700 per condition); weight and opacity scale with $|r|$. Red, positive coupling; blue, anti-correlation. Module assignments from hierarchical clustering (average linkage, Pearson distance $1-r$; cophenetic r=0.804). **(C)** Weighted participation coefficient per region (n=313) ($p < 0.0001$). **(D)** Within-module coupling: mean signed Pearson r from each region to all same-module regions (self-excluded) ($p < 0.0001$). **(E)** Between-module coupling: mean signed Pearson r to all other-module regions ($p < 0.0001$). **(F)** Regions ranked by ΔPC (Inflamed−Control); top 20 shown. NTS, nucleus tractus solitarius; DTN, dorsal tegmental nucleus; LPO, lateral preoptic area; SNr/SNc, substantia nigra; SPV, spinal trigeminal complex; DMH, dorsomedial hypothalamus;

HPF, hippocampal formation; MD, mediodorsal thalamus; RPA, raphe pallidus; STN, subthalamic nucleus; MMl, lateral mammillary nucleus; MEA, medial amygdala; VPL, ventroposterolateral thalamus; LDT, laterodorsal tegmental nucleus; PA, posterior amygdala. **(G)** Inter-module chord diagrams, Control and Inflamed. Ribbon width is proportional to absolute mean Pearson r between module pairs. Red, positive coupling; blue, anti-correlation. *Control: n = 6; Inflamed n = 3. Unpaired t-test: C-E.*

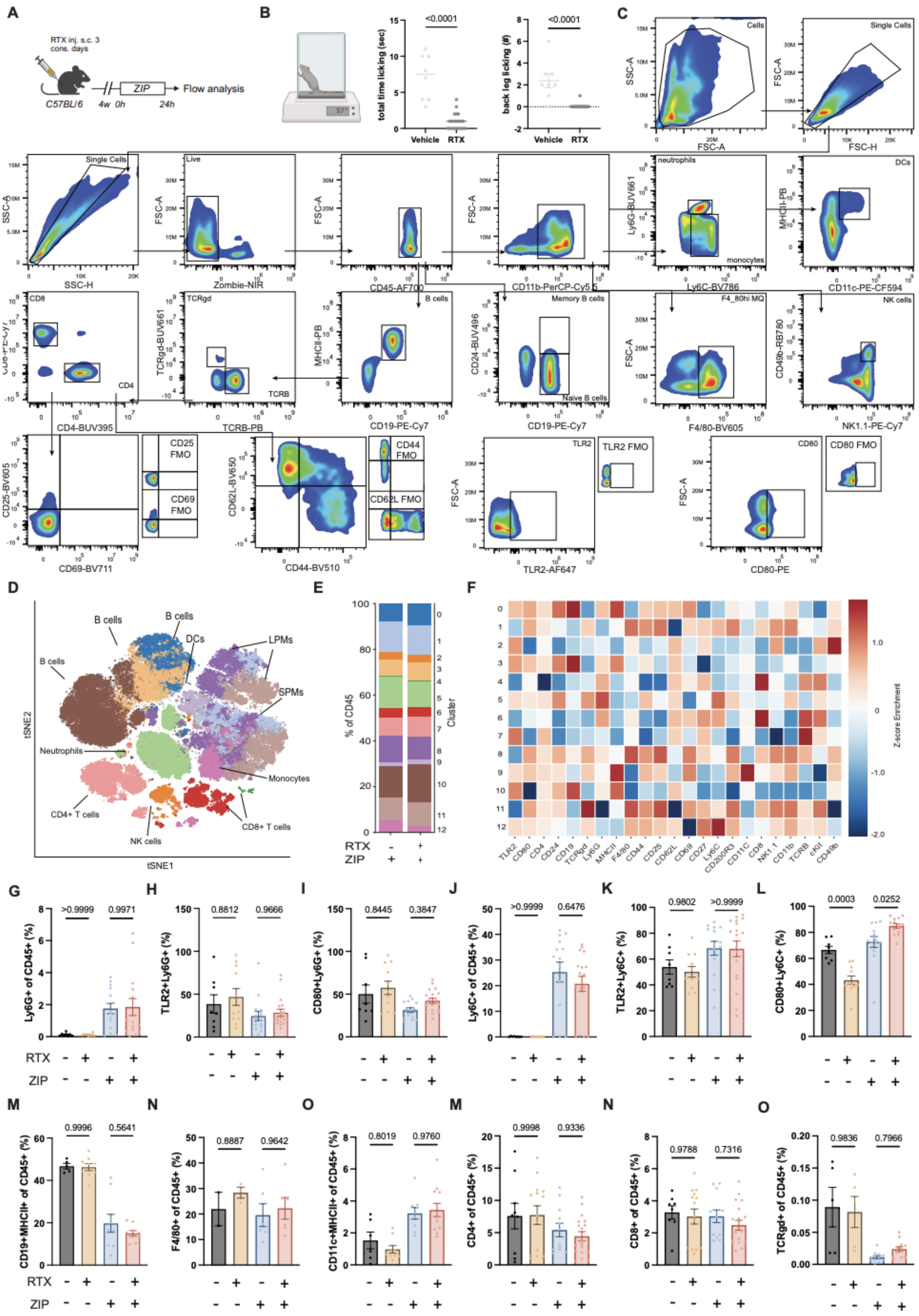

**Fig. S3. Gating strategy and full immune panel for RTX- and zymosan-treated mice.**

**(A)** Experimental timeline for assessing the role of peripheral TRPV1+ neurons in acute Zymosan-induced peritonitis (ZIP). Four groups of 4-week-old C57BL/6 mice received either RTX or vehicle to ablate TRPV1+ neurons. After recovery, one group of RTX treated, and one group of vehicle treated mice were challenged with Zymosan and the other group of RTX treated, and the other group of vehicle treated mice received a vehicle injection. 24 hours later, immune analysis was performed. **(B)** Schematic of hot plate test (left) and measured responses from RTX treated mice. Quantification of the total time (seconds) licking while on the plate (middle) ($p < 0.0001$) and number of licks to the back legs (right) ($p < 0.0001$) Vehicle: n = 8; RTX: n = 23. **(C)** Flow cytometry gating strategy for peripheral immune populations, including FMO controls for CD80, TLR2, CD62L, CD44, CD25, and CD69. **(D)** t-SNE embedding of CD45+ immune cells in the PLF following ZIP in Vehicle (n = 6) vs. RTX (n = 6) pretreated mice. **(E)** Stacked bar charts of the frequency of each cluster out of CD45, per group. **(F)** Heatmap of marker enrichment (z-scored) within each cluster. **(G-O)** Bar plots of the frequency of major cell types in the PLF across each of the 4 groups. Vehicle + Vehicle: n = 8; RTX + Vehicle: n = 11; Vehicle + ZIP: n = 15; RTX + ZIP: n = 18; 3 repeats. *Data presented as mean ± SEM. Statistical significance determined by one-way ANOVA for G-O.*

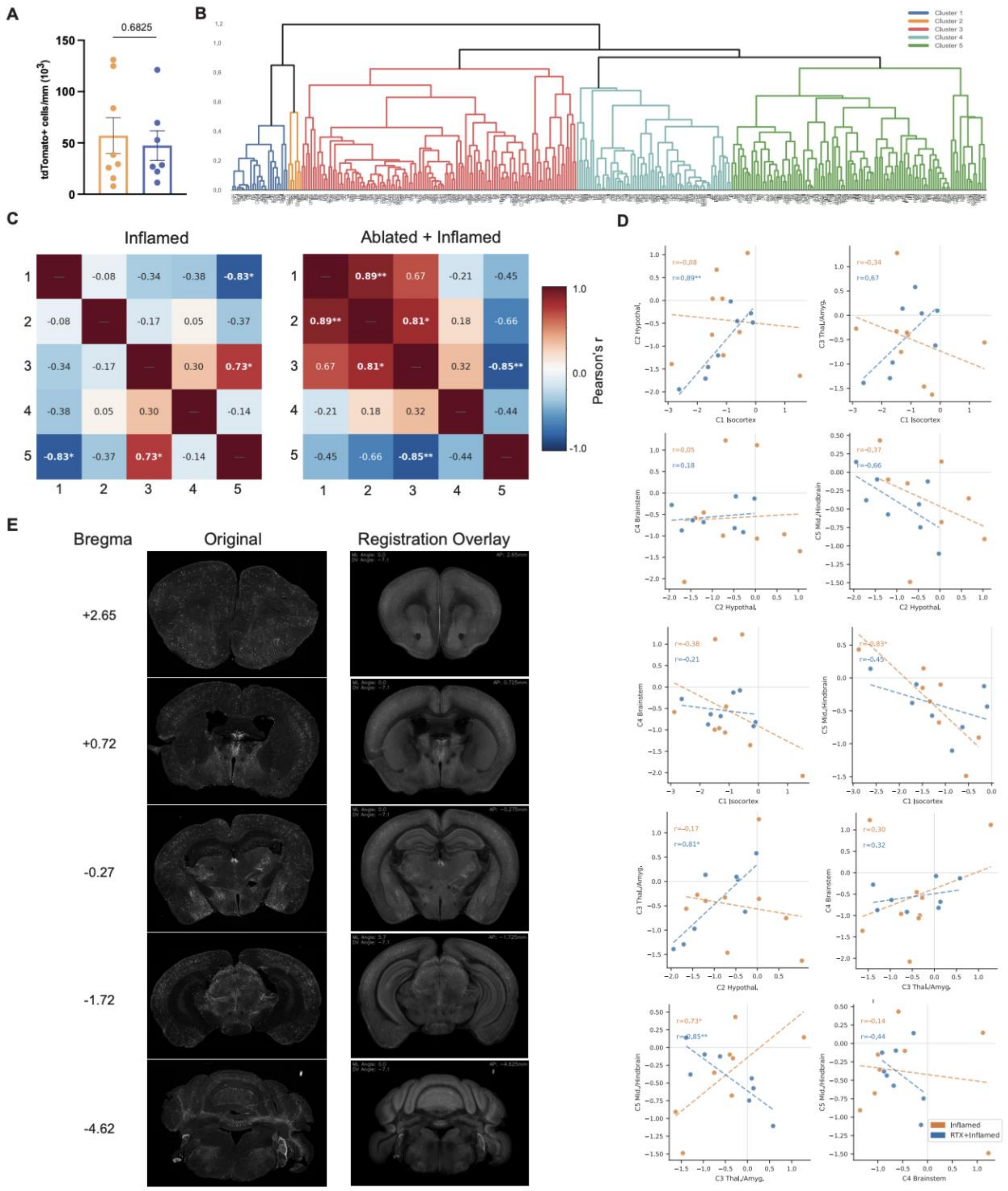


**Figure S4. Nociceptor status restructures inter-cluster covariance without altering total neuronal recruitment.**

**(A)** Total tdTomato+ cell density (cells/mm, ×10³) across the whole brain in Inflamed vs. Ablated + Inflamed mice (p = 0.6825, unpaired t-test). **(B)** Full hierarchical clustering dendrogram of pairwise regional activity correlations (z-scored Pearson r, average linkage; 5,000 permutations) across 361 brain regions. Color-coded cluster assignments (Clusters 1–5) correspond to Fig. 2H. **(C)** Pairwise between-cluster Pearson r correlation matrices for Inflamed (left) and Ablated + Inflamed (right) mice. In Inflamed animals, the isocortex (Cluster 1) and midbrain/hindbrain (Cluster 5) are anti-correlated ($r = -0.83$, $p = 0.011$), while the thalamus/amygdala (Cluster 3) and midbrain/hindbrain (Cluster 5) are positively coupled ($r = 0.73$, $p = 0.021$). After nociceptor ablation, the isocortex–midbrain/hindbrain anti-correlation is lost ($r = -0.45$, $p = 0.263$) and replaced by strong positive covariance between the isocortex, hypothalamus, and thalamus/amygdala ($r_{1\text{–}2} = 0.89$, $p < 0.01$; $r_{2\text{–}3} = 0.81$, $p < 0.05$), while the Cluster 3–5 relationship inverts ($r = -0.85$, $p < 0.01$). *$p < 0.05$, **$p < 0.01$, Fisher's r-to-z test. **(D)** Scatter plots of pairwise inter-cluster activity for Inflamed (orange) and RTX + Inflamed (blue) mice across all ten cluster-pair combinations. Heightened nociception during inflammation shifts inter-cluster coupling patterns relative to controls, particularly strengthening the corticothalamic relationship. Dashed lines, linear regression fit; r, Pearson correlation coefficient. **(E)** Representative whole-brain TRAP2-Ai14d sections at five Bregma levels used for analysis with the DMC plugin (+2.65, +0.72, −0.27, −1.72, −4.62 mm), shown as raw fluorescence and with Allen CCF registration overlay, confirming atlas alignment for region-level quantification. *Inflamed n = 8, RTX+Inflamed n = 9.*

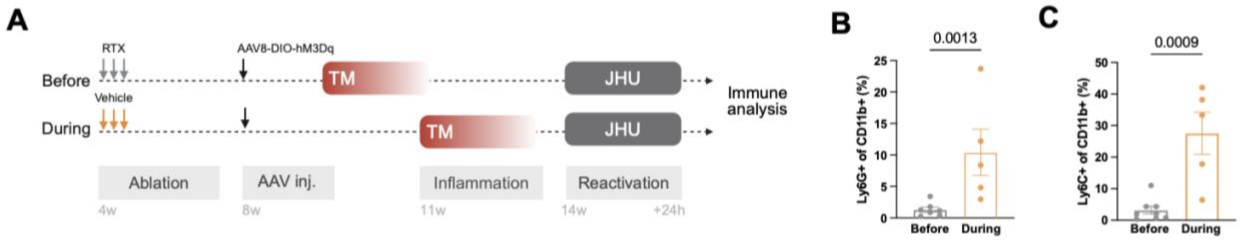


**Figure S5. Validation of pIC reactivation of ZIP in RTX treated mice.**

**(A)** Experimental timeline. Mice were treated with RTX or vehicle to ablate TRPV1+ nociceptors and after recovery, all mice received AAV8-DIO-hM3Dq injected to the pIC. Before group had neurons labeled prior to inflammation and During group had neurons labeled with TM during ZIP. JHU was later injected for chemogenetic reactivation and immune populations were analyzed 24 hours later. **(B)** Bar plot of the frequency of Ly6G+ cells in the PLF after reactivation ($p = 0.0013$). **(C)** Frequency of Ly6C+ after reactivation ($p = 0.0009$). *Data shown as mean ± SEM with individual data points. Unpaired t-tests. Before: n =8, During: n = 5.*

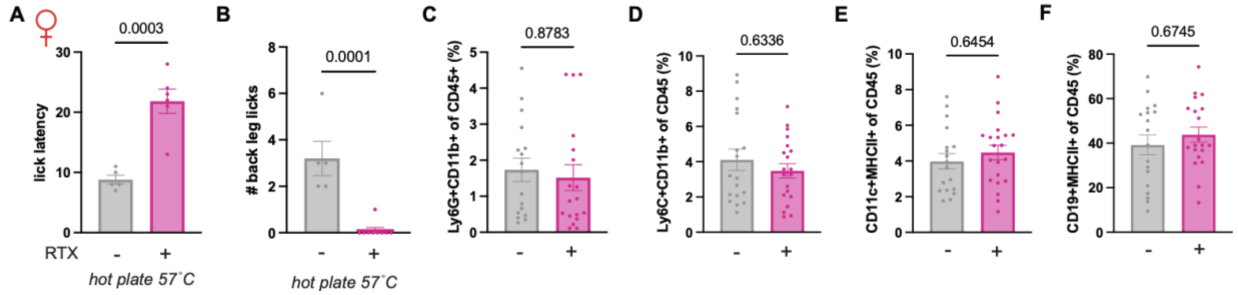


**Figure S6. RTX treated female mice have reduced response to heat but reactivation by pIC neurons is unaffected.**

**(A–B)** Hot plate test (57°C) in female mice. RTX-treated animals had longer lick latency (A; unpaired t-test, p = 0.0003; n = [n] per group) and fewer back leg lick bouts (B; p = 0.0001). **(C–F)** Peripheral immune composition without inflammation: Ly6G+CD11b+ neutrophils (C; p = 0.8783), Ly6C+CD11b+ monocytes (D; p = 0.6336), CD11c+MHCII+ dendritic cells (E; p = 0.6454), and CD19+MHCII+ B cells (F; p = 0.6745) of CD45+ cells were comparable between RTX − and RTX + groups (unpaired t-test; n = [n] per group). *Mean ± SEM with individual data points.*

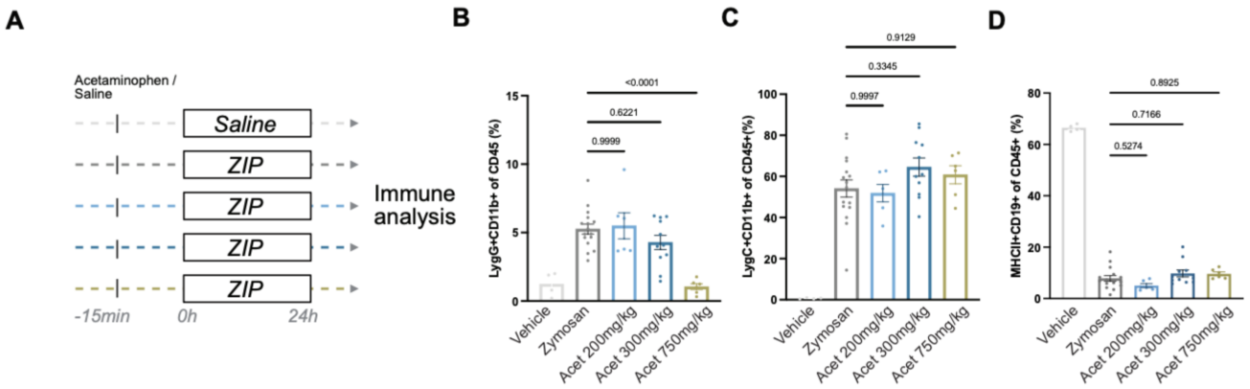


**Figure S7. Acetaminophen suppresses neutrophil recruitment during ZIP only at the highest dose.**

**(A)** Experimental design. Acetaminophen (200, 300, or 750 mg/kg) or saline was given 15 minutes before intraplantar zymosan (ZIP); immune populations analyzed at 24 hours. **(B)** Ly6G+CD11b+ neutrophil frequency of CD45+. Only 750 mg/kg reduced neutrophil recruitment relative to zymosan alone (one-way ANOVA with Tukey's; Vehicle vs. Zymosan: $p = 0.9999$; Zymosan vs. Acet 200 mg/kg: $p = 0.6221$; Zymosan vs. Acet 750 mg/kg: $p < 0.0001$; n = [n] per group). **(C)** Ly6C+CD11b+ monocyte frequency of CD45+; no significant differences (Zymosan vs. Acet 300 mg/kg: $p = 0.3345$; vs. Acet 750 mg/kg: $p = 0.9129$). **(D)** MHCII+CD19+ B cell frequency of CD45+; unchanged across doses (Acet 200: $p = 0.5274$; Acet 300: $p = 0.7166$; Acet 750: $p = 0.8925$). *Mean ± SEM with individual data points.*

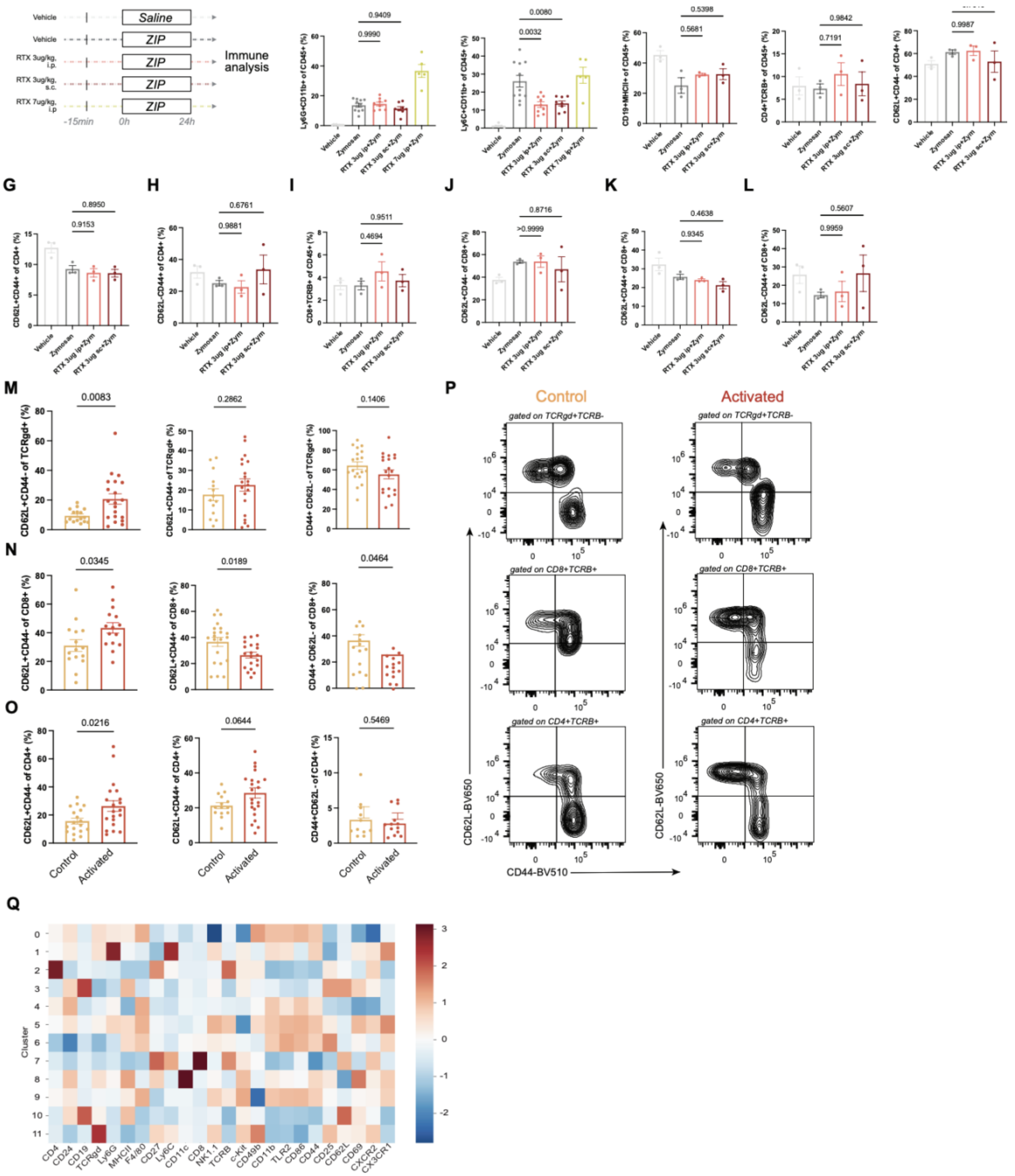


**Fig. S8 RTX amplifies zymosan-induced myeloid recruitment in a dose- and route-dependent manner; T cell populations are unaffected.**

**(A)** Timeline. RTX (3 μg/kg i.p., 3 μg/kg s.c., or 7 μg/kg i.p.) or vehicle was given 15 minutes before intraplantar zymosan (ZIP); peripheral immune populations analyzed at 24 hours. **(B)** Ly6G+CD11b+ neutrophil frequency of CD45+. RTX at 7 μg/kg i.p. significantly elevated neutrophils relative to zymosan (one-way ANOVA with Tukey's; Veh vs. Zym: $p = 0.9990$; Zym vs. RTX 3 µg/kg i.p.: $p = 0.9409$; Zym vs. RTX 7 µg/kg i.p.: $p < 0.0001$; n = [n] per group). **(C)** Ly6C+CD11b+ monocyte frequency of CD45+. Both RTX 3 µg/kg i.p. and 7 µg/kg i.p. elevated monocytes compared to zymosan alone (Zym vs. RTX 3 µg/kg i.p.: $p = 0.0032$; vs. RTX 7 µg/kg i.p.: $p = 0.0080$; RTX 3 µg/kg s.c. vs. RTX 7 µg/kg i.p.: $p = 0.9526$). **(D)** CD19+MHCII+ B cells of CD45+; no changes across groups (all $p > 0.5$). **(E)** CD4+TCRB+ T cells of CD45+; unchanged (all $p > 0.7$). **(F–H)** CD4+ memory subsets: CD62L+CD44+ effector memory (F; all $p > 0.7$), CD62L+CD44+ (G; $p = 0.9153$, 0.8950), and CD62L−CD44+ (H; $p = 0.9881$, 0.6761) CD4+ T cells were unchanged. **(I–L)** CD8+ subsets: CD8+TCRB+ (I; $p = 0.4694$, 0.9511), CD62L+CD44+ CD8+ (J; $p > 0.9999$, 0.8716), CD62L+CD44+ CD8+ (K; $p = 0.9345$, 0.4638), and CD62L−CD44+ CD8+ (L; $p = 0.9959$, 0.5607) (**M)** γδ T cell analysis: CD62L+CD44− of TCRγδ + ($p = 0.0083$); CD62L+CD44+ of TCRγδ + ($p = 0.2862$); CD44+CD62L− of TCRγδ + ($p = 0.1406$). **(N)** CD8+ T cell analysis: CD62L+CD44− of CD8+ ($p = 0.0345$); CD62L+CD44+ of CD8+ ($p = 0.0189$); CD44+CD62L− of CD8+ ($p = 0.0464$). **(O)** CD4+ T cell analysis: CD62L+CD44− of CD4+ ($p = 0.0216$); CD62L+CD44+ of CD4+ ($p = 0.0644$); CD44+CD62L− of CD4+ ($p = 0.5469$). **(P)** Representative flow cytometry plots showing CD62L vs. CD44 gating on TCRγδ + (top), CD8+ (middle), and CD4+ (bottom) T cells for Control (left) and Activated (right) animals. **(Q)** Heatmap of marker enrichment (z-scored) within each cluster. *(B-L: one-way ANOVA with Tukey's) (M-O: unpaired t-test). Mean ± SEM with individual data points.*
